# Microtubule-based nucleation results in a large sensitivity to cell geometry of the plant cortical array

**DOI:** 10.1101/2024.03.25.586463

**Authors:** Marco Saltini, Eva E. Deinum

## Abstract

Many plant cell functions, including cell morphogenesis and anisotropic growth, rely on the self-organisation of cortical microtubules into aligned arrays with the correct orientation. An important ongoing debate is how cell geometry, wall mechanical stresses, and other internal and external cues are integrated to determine the orientation of the cortical array. Here, we demonstrate that microtubule-based nucleation can markedly shift the balance between these often competing directional cues. For this, we developed a novel, more realistic model for microtubule-based nucleation in the simulation platform CorticalSim, which avoids the longstanding inhomogeneity problem stemming from previous, less realistic models for microtubule-based nucleation. We show that microtubule-based nucleation increases the sensitivity of the array to cell geometry, and extends the regime of spontaneous alignment compared to isotropic nucleation. In the case of cylindrical cell shapes, we show that this translates into a strong tendency to align in the transverse direction rather than along the vertical axis, and this is robust against small directional cues favouring the longitudinal direction. Comparing various cylinders and boxes, we show that different nucleation mechanisms result in different preferred array orientations, with the largest differences on cylinders. Our model provides a powerful tool for investigating how plant cells integrate multiple biases to orient their cortical arrays, offering new insights into the biophysical mechanisms underlying cell shape and growth.

## Introduction

Plants take their shape through coordinated processes of cell division, expansion and differentiation. At the cellular level, many of these processes are shaped by the plant cortical microtubule array [1–3]. This dynamic structure of membrane associated microtubules guides the anisotropic deposition of cellulose microfibrils, the main load bearing component of the cell wall, and predicts the future cell division site [4–7]. Under the right conditions [8, 9], these dynamic microtubules can self-organise into a highly aligned array. This self-organisation depends on frequent collisional interactions among microtubules, resulting in angle-dependent outcomes [10]. The occurrence of such collisional interactions is controlled by the dynamics of individual microtubules which, hence, play a crucial role in the self-organisation of the array [8, 11]. Microtubule nucleation, which predominantly occurs from exiting microtubules (Figure 1A; [12, 13]), has a profound impact on the self-organisation of the cortical microtubule array and its ability to adopt complex patterns [13–18]. The degree of co-alignment between new and parent microtubules is an important factor in explaining these effects (Figure 1A) [17, 18]. Consequently, any quantitative predictions from computer simulations critically depend on a sufficiently realistic implementation of microtubule nucleation.

**Figure 1:**
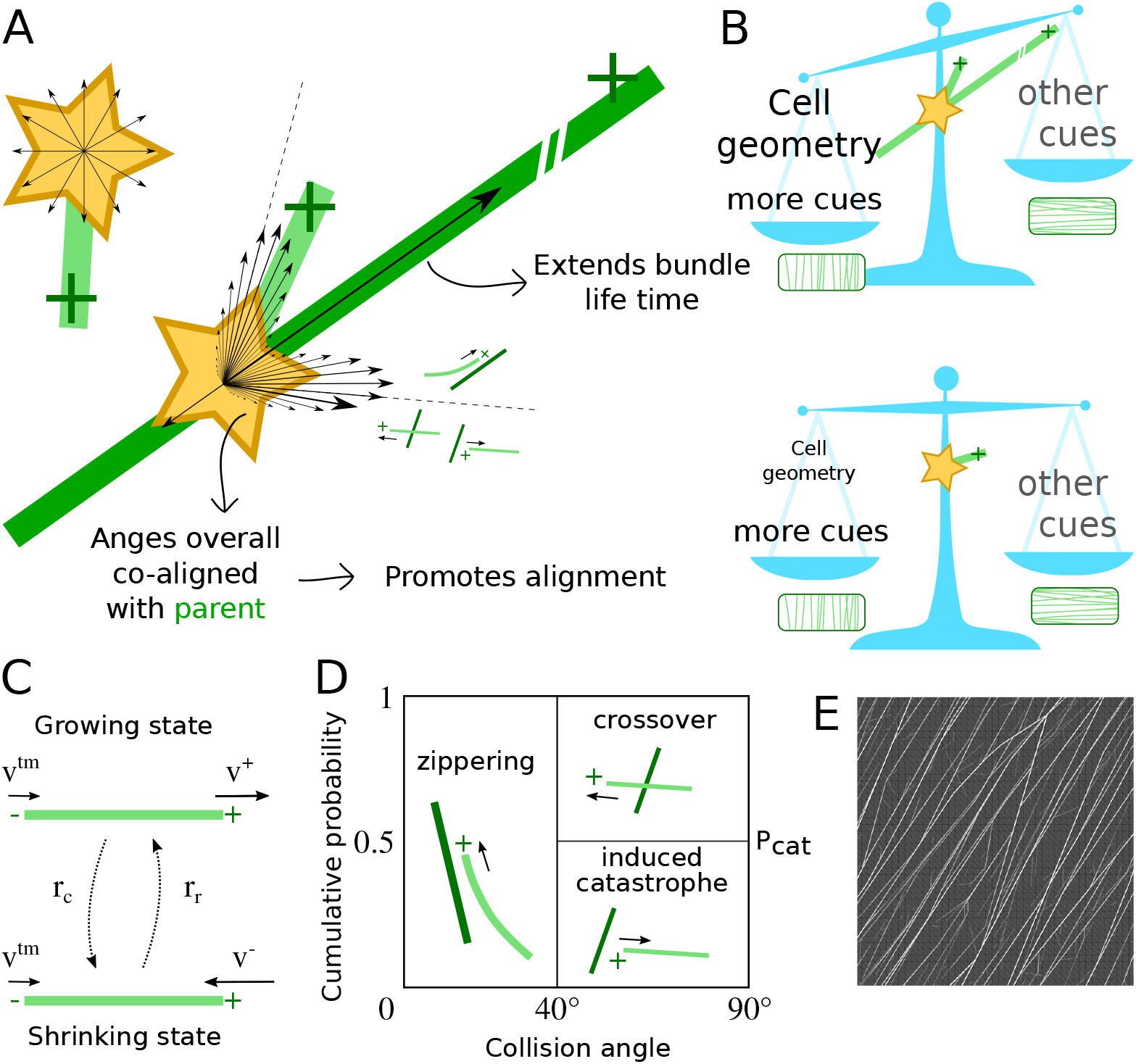
Different nucleation modes (and their hypothetical impact). (A) Unbound nucleation (i.e., nucleation occurring from a nucleation complex dispersed in the cytoplasm) typically has a uniformly distributed orientation, whereas microtubule-based nucleation follows a specific distribution relative to the parent microtubule, with parallel, antiparallel and “branched” components. The part parallel to the parent microtubule (31% in our simulations [17]) can increase the life time of a microtubule bundle. Additionally, the distribution of nucleation angles is overall co-aligned with the parent microtubule, which increases the parameter regime of spontaneous microtubule alignment. Dashed lines indicate the range of zippering onto microtubules with the same orientation as the parent. The arrow with the large arrowhead (one side only) indicates a 45^°^ angle, i.e., neutral with respect to co-alignment with the parent. (B) Working hypothesis: as a consequence of the properties of microtubule-based nucleation, the persistence of bundles increases, which allows for longer retention and further communication of the information about the array stored in that bundle. This renders the array as a whole more sensitive to global cues like the size and shape (geometry) of the cell. The impact of different cues varies, as visualized by variation in font size. (C-E) **Overview of model dynamics**. (C) Dynamics of individual microtubules. The plus-end (+) stochastically switches between the growing and shrinking state, whereas the minus-end (-) shows steady retraction. (D) Angle dependent collision outcomes. (E) Fragment of an example simulated array. Individual microtubules are drawn with a degree of transparency, so thicker bundles appear as stronger white lines.

This is particularly relevant in the ongoing debate about what controls the orientation of the cortical microtubule array (Figure 1B), which is the focus of this manuscript. Array orientation is significantly influenced by a combination of global and local cues. In this manuscript, we consider *global cues* as cues that require information processing on the whole cell to respond to them, while *local cues* can be responded to with only local information. For example, global cues such as the cell geometry as a whole can favour array orientation along the shortest closed circumferences of the cell [19, 20]. Local cues, including the alignment of microtubules with mechanical stresses [21], could select for a specific orientation among these, or potentially even override the global cues [22, 23]. Given that new microtubules in plants typically nucleate in directions strongly correlated with the orientation of their parent microtubule [12, 17, 18], how microtubules are nucleated in simulations of the cortical array is crucial. Indeed, different nucleation modes result in different sensitivities to local and global cues [14, 15, 24, 25]. It is hypothesised that the basis of this different behaviour is that microtubule-based nucleation increases bundle life time and, thereby, longer maintains the information stored in the constituent microtubules [17, 24]. Therefore, the ability to realistically simulate microtubule nucleation is critical for any computational study that involves a competition between local and global cues affecting alignment and orientation.

Explicit stochastic simulations (Figure 1C-E) have a long tradition in the study of cortical array behaviour (e.g., see [9, 24] for reviews). Despite the importance of microtubule-based nucleation, however, most computational and theoretical research on microtubule arrays has traditionally employed isotropic nucleation, where microtubules are initiated at random positions with random orientations. While helpful for understanding the effects of microtubule collisions [8, 10, 26] and geometrical constraints on alignment [19, 22, 27–29], this approach ignores the fact that microtubules often nucleate from existing ones [12, 13, 30, 31]. Many simulation studies with microtubule-based nucleation [15, 17, 32, 33], however, suffer from a so-called “inhomogeneity problem” [14]. In essence, this is a computational problem arising from a naive implementation of microtubule-based nucleation, i.e., when the local share of microtubule-based nucleation is *linearly* dependent on the local density. This linear density-dependence causes an unrestricted positive feedback on local microtubule density, leading to overly dense regions and very sparse areas in between. As cellulose synthase complexes are preferentially inserted near microtubules [6], such inhomogeneities would lead to structural weaknesses in the cell wall. While this feedback loop originates from the biological reality of microtubule-based nucleation, the naive implementation has no ingredient counteracting the positive feedback. In reality, however, the local nucleation rate saturates with density, thereby ensuring homogeneity [14].

Jacobs et al. [14] introduced a nucleation mechanism that naturally creates this saturating local density-dependence by explicitly considering appearance and diffusion of nucleation complexes at the membrane. Applied to spatially distributed but non-interacting microtubules^1^, this mechanism resulted in homogeneous arrays. In those simulations, however, nucleation complex diffusion takes up a large fraction of the computational time, even though nucleation itself is a relatively rare event. The diffusion of explicit nucleation complexes is also employed in CytoSim (e.g., [34]), which allows for microtubule-microtubule interactions, but is overall computationally heavy. The event-based platform CorticalSim [11], in contrast, has been developed for high speed, which allows for accurate computation of ensemble statistics from many simulations combined with exploration of parameter space in relatively short time [5, 11, 15, 17, 19, 35], exactly what is needed for the quantitative investigation of competing cues. Until now, however, this and all other relatively fast platforms lacked a nucleation algorithm along the lines of Jacobs et al. [14]^2^.

Here, we bring the possibility of local density-dependent nucleation (LDD) to the context of interacting cortical microtubules: we describe a new algorithm in CorticalSim that efficiently approximates the diffusion of nucleation complexes. This algorithm overcomes the long-standing issue of inhomogeneity at limited computational cost. We compare arrays produced by LDD nucleation mode with those generated by global density-dependent microtubule-based nucleation (GDD), and isotropic nucleation (ISO). For nucleation, “local” means that the local nucleation rate and the probability that a specific spot attracts a particular nucleation event are fully determined by the density and array structure in a small area (in our case, a circle with a few *μ*m radius). “Global” means that every unit of length of a microtubule competes with equal weight for a particular nucleation event, whether it is in a thick bundle of alone. We show that cell geometry has a much stronger impact on array orientation than previously thought. More generally, our results show that different nucleation mechanisms have major impacts in the responsiveness of the array to cues of different types.

## 1 Results

### 1.1 Microtubule-based nucleation supports homogeneous arrays

The hypothesis that microtubule-based nucleation improves alignment of the cortical array through extending bundle life time [17] is critical to understanding the impact of this type of nucleation. Previously published results including microtubule-based nucleation, however, all depended on a mechanism that introduces a global competition for nucleation (global density-dependence (GDD) in our study), so it cannot be ruled out that some of the observed behaviour was caused by the inhomogeneity problem rather than microtubule-based nucleation *per se*. We first verified that our algorithm for local density dependence (LDD) produces homogeneous arrays on both planar and capped cylindrical surfaces. Figure 2 shows that LDD indeed produced homogeneous arrays with default parameters (Table 1), comparable to isotropic nucleation (ISO), whereas GDD resulted in large inhomogeneities, visible as a combination of both large, nearly empty areas and a few very dense bundles. With LDD and ISO, inhomogeneities in the early array disappeared over time, whereas with GDD, array inhomogeneity increased with time (Figure 2A-E). We further quantified the development of inhomogeneity by following the local density over time in transversely oriented arrays on cylinders (Figure 2F). Total density and its local variations were similar with ISO and LDD. With GDD, however, all density tended to accumulate in a single transverse band. Once established, such bands were extremely stable and reached unrealistically high densities (saturating at values over 120 *μ*m^−1^, equivalent to 3 layers of microtubules, touching side by side, on top of each other). The local density profiles with LDD and ISO, moreover, were more dynamic than with GDD. Such fluctuations in microtubule locations could further support homogeneous cell wall properties (see also [14]).

**Table 1:** Default model parameters.

| Parameter | Description | Value | Unit | Source |
| --- | --- | --- | --- | --- |
| $v^+$ | Growth speed | 0.05 | $\mu\text{m s}^{-1}$ | [15] |
| $v^-$ | Shrinkage speed | 0.08 | $\mu\text{m s}^{-1}$ | [15] |
| $v^{\text{tm}}$ | Treadmilling speed | 0.01 | $\mu\text{m s}^{-1}$ | [17, 59] |
| $r_c$ | Catastrophe rate | variable | $\text{s}^{-1}$ | |
| $r_r$ | Rescue rate | 0.001 | $\text{s}^{-1}$ | [15] |
| $\theta_c$ | Maximum collisional angle for bundling | $40^\circ$ | - | [10] |
| $P_{\text{cat}}$ | Fraction of induced catastrophe after collision when $\theta > \theta_c$ | 0.5 | - | [11, 35] |
| $r_n$ | Nucleation rate for ISO and GDD | 0.001 | $\mu\text{m}^{-2} \text{s}^{-1}$ | [15] |
| $r_{\text{ins}}$ | Complex appearance rate for LDD | 0.0045 | $\mu\text{m}^{-2} \text{s}^{-1}$ | [14] |
| $r'_{\text{ins}}$ | rescaled: | 0.00108 | $\mu\text{m}^{-2} \text{s}^{-1}$ | |
| $r_u$ | Per-complex unbound nucleation rate | 0.002 | $\text{s}^{-1}$ | [14] |
| $D$ | Diffusion coefficient of nucleation complexes | 0.013 | $\mu\text{m}^2 \text{s}^{-1}$ | [14] |
| $N$ | Number of meta-trajectories for LDD | 6 | - | |
| $R$ | Maximum meta-trajectory length for LDD | 1.5 | $\mu\text{m}$ | |
| $P_{\text{rej},b}$ | Rejection probability for microtubule-based nucleation for LDD | 0.76 | - | [13, 14] |
| $P'_{\text{rej},b}$ | rescaled: | 0 | - | |
| $P_{\text{rej},u}$ | Rejection probability for unbound nucleation for LDD | 0.98 | - | [14] |
| $P'_{\text{rej},u}$ | rescaled: | 0.91667 | - | |
| $\rho_{1/2}$ | Tubulin density at which half of nucleations are isotropic for GDD | 0.1 | $\mu\text{m}^{-1}$ | [17] |
| $f_{\text{forward}}$ | Nucleation probability along the parent for GDD and LDD | 0.31 | - | [17] |
| $f_{\text{backward}}$ | Nucleation probability anti-parallel to the parent for GDD and LDD | 0.07 | - | [17] |
| $f_{\text{left}}$ | Nucleation probability to the left of the parent for GDD and LDD | 0.31 | - | [17] |
| $f_{\text{right}}$ | Nucleation probability to the right of the parent for GDD and LDD | 0.31 | - | [17] |
| $\epsilon$ | Ellipse eccentricity for GDD and LDD | 0.89 | - | [12, 17] |
| $\theta_b$ | Mean branching angle for GDD and LDD | $35^\circ$ | - | [12, 17] |

**Figure 2:**
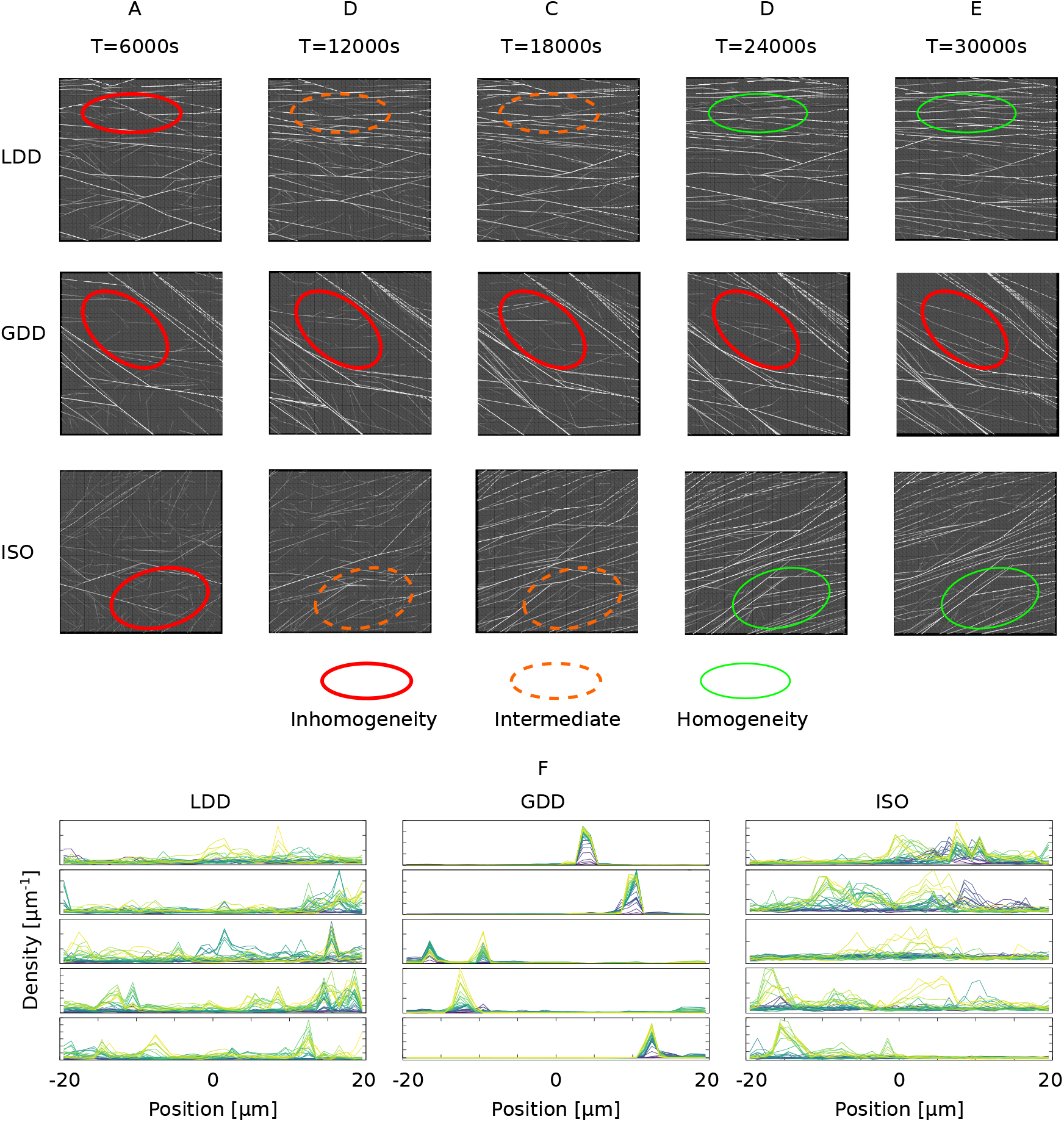
Simulation snapshots show that the new LDD microtubule-based nucleation yields much more homogeneous arrays than GDD microtubule-based nucleation. (A-G) Simulation snapshots of a 40 × 40*μ*m^2^ square domain taken every 6000 s (100 min). Red thick ovals represent areas of inhomogeneity, orange dashed ovals intermediate areas, and green thin ovals areas where the array became homogeneous. Note the persistence of areas of inhomogeneity in the GDD case, compared to the transient inhomogeneity in the LDD and ISO cases. (F) Local density along the axis of a cylinder (L=40 *μ*m; R=6 *μ*m) in bins of 1 *μ*m for tightly transversely oriented arrays. Five independent simulations per nucleation mode. Time increases from purple (start of simulation) to yellow (end). In these simulations, catastrophe rate values correspond to data points marked with large symbols in Figure 3, i.e., (LDD) *r*_*c*_ = 0.00275 s^−1^, (GDD) *r*_*c*_ = 0.003 s^−1^, (ISO) *r*_*c*_ = 0.002 s^−1^. Density values are omitted from the axes for readability. Tic marks on the vertical axis represent density increments of 3 *μ*m^−1^ for LDD and ISO and 40 *μ*m^−1^ for GDD. All axes start at 0. Transverse arrays were randomly selected from a larger set of simulations with the sole criterium that the array orientation angle on the cylinder mantle 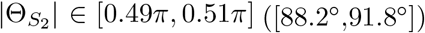.

### 1.2 Microtubule-based nucleation stimulates array alignment

Although with LDD the visual appearance was rather similar to arrays produced with isotropic nucleation, LDD microtubule-based nucleation increased the parameter regime for spontaneous alignment almost as much as the GDD algorithm (Figure 3A). In this figure, the parameters describing the dynamic instability of individual microtubules and the nucleation rate are collapsed onto a single number *G*, defined in Equation (8) [8, 17, 36]. This so-called control parameter is the (negative) ratio of two length scales: an interaction length scale and the average microtubule length without interactions. As *G* increases towards zero, the average number of interactions per microtubule life time increases. With sufficient interactions, spontaneous alignment occurs (order parameter *S*_2_ on the vertical axis increases from near zero to near one. See Equation (6) for a definition of *S*_2_). With this understanding of the process [8, 9, 20, 24], the result in Figure 3A can be restated as follows: microtubule-based nucleation decreases the minimal number of interactions per average microtubule life time that is required for spontaneous alignment (other parameters remaining constant). Note that in the LDD case, *G* is a measured quantity that depends on the actual frequency of nucleation events (Equation (9)), which is inherently stochastic. However, as shown in Figure S1B, the error bars of the measured *G* were small, as expected from the slow scaling of *G* with 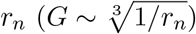.

**Figure 3:**
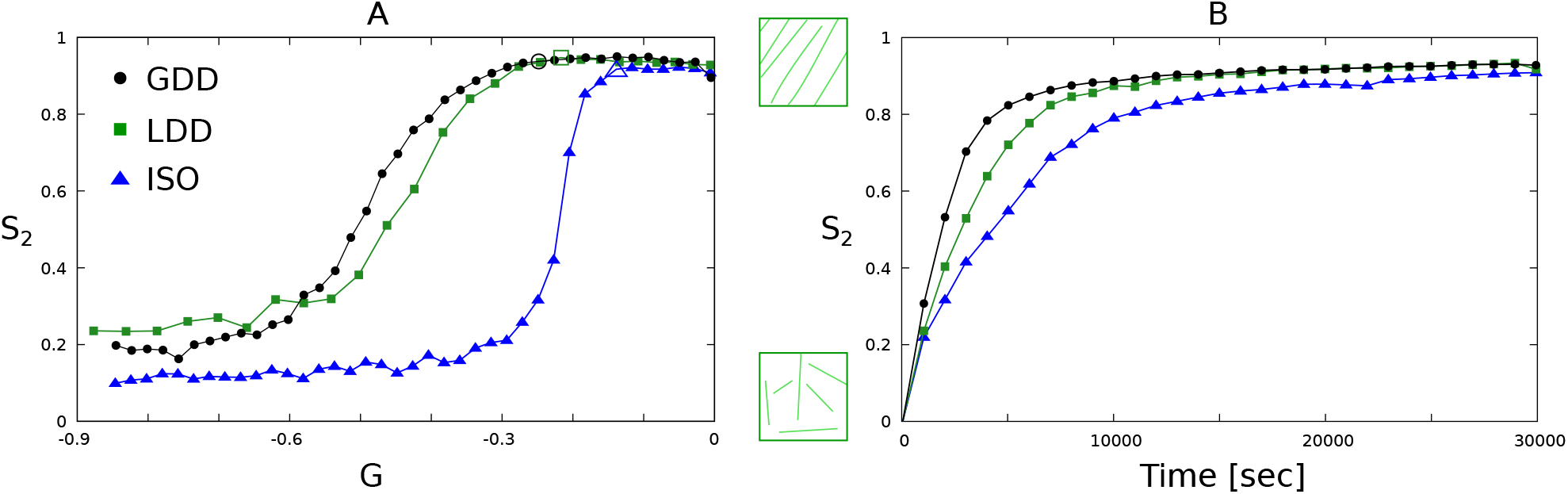
LDD nucleation maintains the alignment promoting effect of microtubule-based nucleation as previously reported for GDD. Aligned regime for LDD, GDD, and ISO nucleation modes on a 40×40 *μ*m^2^ square periodic geometry. (A) Median over 100 independent simulation runs of the order parameter *S*_2_ as function of the control parameter *G* at *T* = 3 × 10^4^ s (8 h, 20 min). Data points marked as larger, empty circle, square, or triangle correspond to the parameter values used for panel (B). (B) Median over 100 independent simulation runs of the order parameter *S*_2_ as function of the simulation time for selected values of *G* (data points marked with large symbols in panel (A)) in the aligned regime. Error bars of (A) can be found in Figure S1.

Alignment also occurred faster with microtubule-based than with isotropic nucleation, when comparing parameters at similar points in the *S*_2_(*G*)-curve (Figure 3B). However, LDD took somehow longer to align than GDD. We expected that the difference between LDD and GDD most resulted from the improved realism of the LDD algorithm: LDD incorporates the experimental observation that nucleation complexes nucleate with higher rate when they are attached to microtubules [13, 15], whereas GDD uses the same rate for bound and unbound complexes. Consequently, the initial nucleation rate is reduced with LDD, but not with the other algorithms. We, therefore, also computed the curves with reduced unbound nucleation rate for GDD and with equal rates for bound and unbound complexes in LDD. Interestingly, the reduction of the unbound rate lead to larger aligned regime for both LDD and GDD (Figure S2A,B). Also with both LDD and GDD, the “noise level” *S*_2_ in the disordered regime increased with unequal rates. With LDD, alignment also occurred slower with equal rates than with reduced unbound rate (Figure S2C). In retrospect, an explanation for this is that, with equal rates, there is more isotropic nucleation and, therefore, the behaviour is more similar to isotropic. With GDD, the density often increased so rapidly (e.g., see Figure 2F), that the fraction of isotropic nucleation had little impact on the time course. So, microtubule-based nucleation speeds up alignment even when the initial nucleation rate is reduced (and see [14, 37]).

Together, these results validate the conclusion that microtubule-based nucleation promotes alignment. Note that this will still depend on the distribution of nucleation angles relative to the parent microtubule (see [17, 18]), which is experimentally observed to be (moderately) co-aligned with the parent microtubule [12, 13].

### 1.3 Microtubule-based nucleation increases sensitivity to geometrical cues

The consequence of increased microtubule bundle lifetime is that the information about the array stored in the bundle is maintained longer. Microtubule-based nucleation could moreover facilitate increased bundle length, which allows information to propagate further through the array. Both processes could change the sensitivity of the array to geometric cues and shift the balance between different types of cues (Figure 1B). This would have major implications for the ongoing debate on what determines the orientation of the cortical array [5, 24, 28, 38, 39]. We, therefore, investigated the preferred orientation of simulated arrays with the three nucleation modes on cylindrical cells of the same aspect ratio but different size. We studied the case of cylinders with 40 *μ*m length and 12 *μ*m diameter, and 60 *μ*m length and 18 *μ*m diameter, which is in the realistic range for epidermal cells of the root and hypocotyl of *Arabidopsis thaliana*. Figure 4 shows that with LDD and ISO, the cylindrical geometry favoured a transverse orientation over a longitudinal one. A high degree of alignment was obtained faster for transverse than for longitudinal arrays (Figure 5). Both effects were stronger for LDD than for ISO.

**Figure 4:**
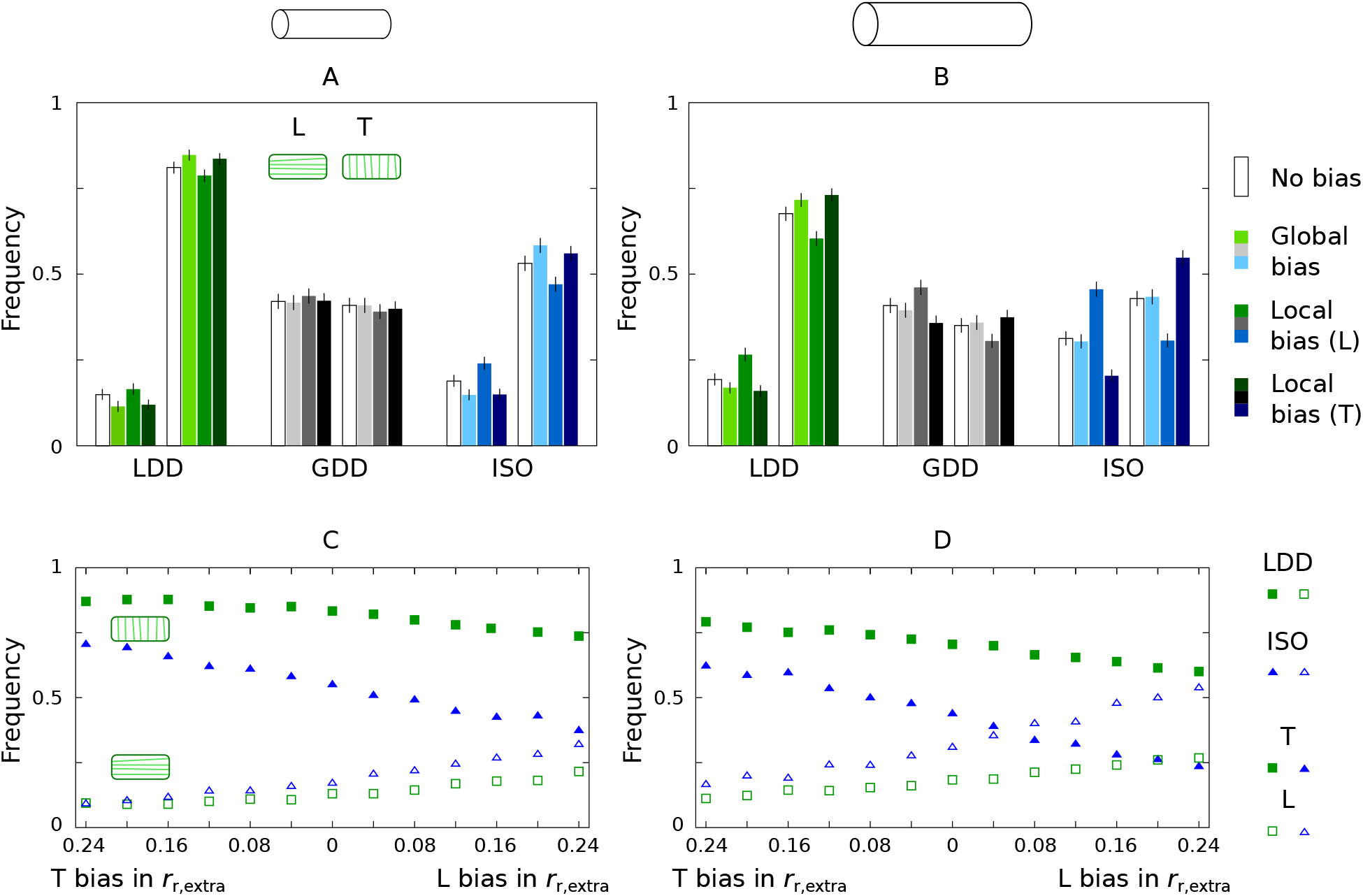
LDD microtubule-based nucleation increases the sensitivity of array orientation to cell geometry. Number of aligned arrays at *T* = 3 × 10^4^ s (8 h, 20 min) with a transverse (T, histograms to the right or filled symbols; when |Θ_2_| *<* 10^°^), or longitudinal (L, histograms to the left or empty symbols; when |Θ_2_| *>* 80^°^) orientation out of n=2000 independent runs each, with or without directional biases: (A,B) 8% increase in catastrophe rate at the cylinder caps for the global bias; and 8% increase in maximum rescue rate on the cylinder mantle according to Equation (10) for the local bias, and (C,D) for different values of the local bias in *r*_*r*,extra_ according to Equation (10); (C,D): in the left half of each panel, the local bias is implemented in the transverse direction, in the right half in the longitudinal direction. Note that, with these parameters, a bias of 0.04 in *r*_*r*,extra_ corresponds to a change of 0.01 in *G* in the same direction. The simulation domain is a cylinder of (A,C) 40 *μ*m length and 12*μ*m diameter, and (B,D) 60 *μ*m length and 18 *μ*m diameter. (A,B) Error bars represent 95% confidence interval according to a binomial model (computed using the binom.test function in R statistical package version 3.6.3). (C,D) Error bars have been omitted because they are approximately the same size as the symbols. In these simulations, *r*_*c*_ = 0.00225 s^−1^.

**Figure 5:**
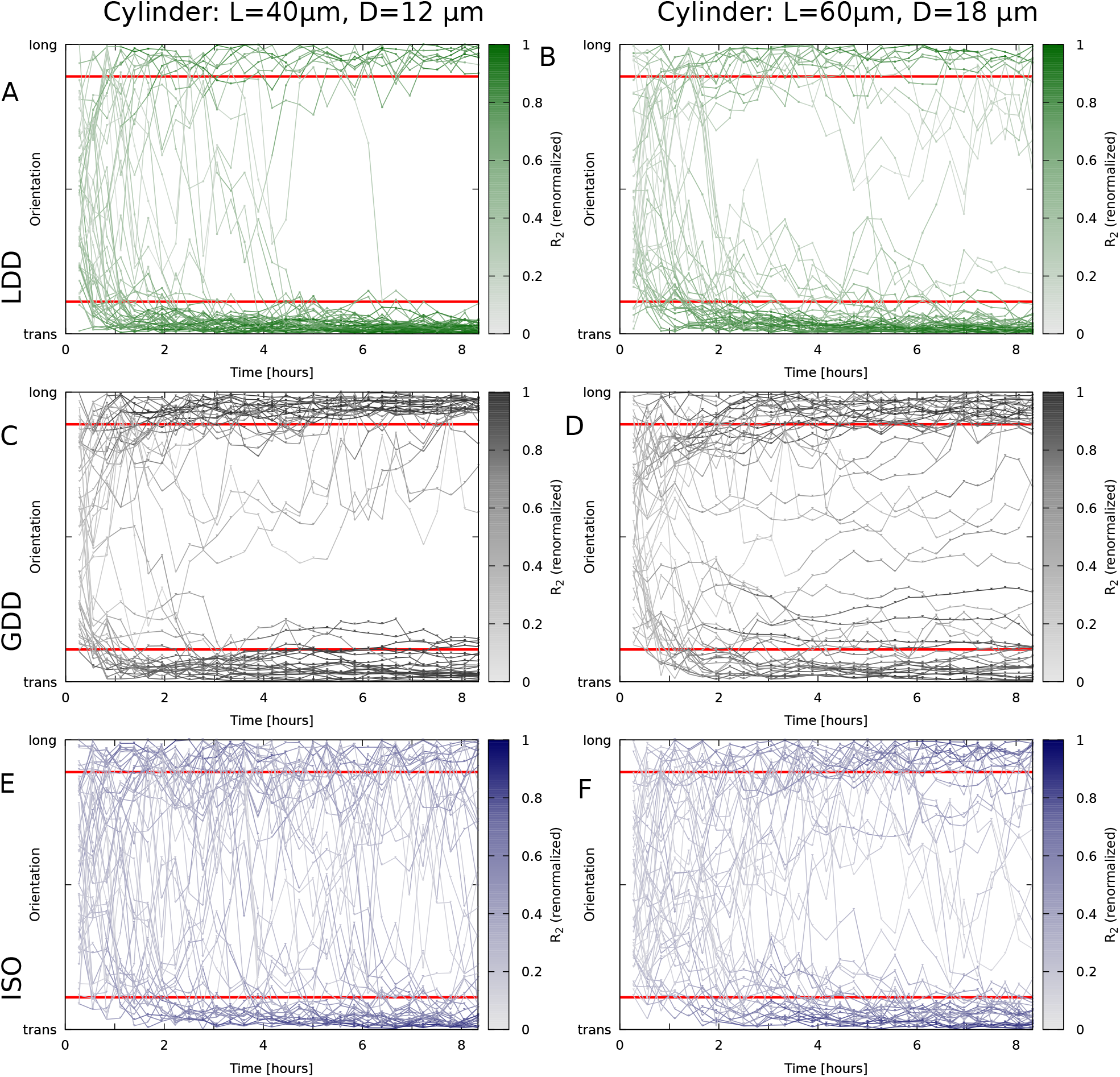
Orientation over time with for individual simulations. Lines depict the orientation of the array (Θ based on *R*_2_) for n=50 simulations per nucleation mode / cylinder size. Lines are coloured by the degree of alignment, renormalized *R*_2_ = *R*_2_*/R*_2_(*max*|Θ). Red lines indicate the boundaries of the “transverse” (trans) and “longitudinal” (long) categories in Figure 4. Microtubule dynamic parameters were the same everywhere in the domain, regardless of orientation. Nucleation modes: (A,B) LDD; (C,D) GDD; (E,F) ISO. Cylinder sizes: length 40 *μ*m, diameter 12 *μ*m (A,C,E); length 60 *μ*m, diameter 18 *μ*m (B,D,F).

GDD, however, behaved differently, at first sight appearing almost indifferent about transverse / longitudinal. As we discuss later, however, the situation is more complicated, and GDD can also be the most sensitive to particular geometrical features.

Both increasing the cell size and switching from LDD to isotropic nucleation increased the sensitivity of array orientation to additional cues that either reinforce or compete with the effect of the geometry alone (Figure 4B). In other words: LDD increased the sensitivity to the global geometry cue. The impact of a higher catastrophe rate at the cylinder caps (“global bias: an 8% increase of *r*_*c*_ at the caps) was weaker on the larger cells. In contrast, the impact of local cues (“local bias”, an 8% increase of rescue rate *r*_*r*_ in the indicated direction), was stronger on the larger cells. As an 8% increase in *r*_*r*_ for the local bias only resulted in a 2% increase in *G*, we varied the strength of the local bias over a larger range (Figure 4C,D, S3). In particular, even with isotropic nucleation, the largest bias tested towards longitudinal still resulted in a small majority of transverse arrays on the small cylinders (maximum 24% increase of *r*_*r*_, corresponding to a 6% increase of *G* for perfectly longitudinal microtubules).

The different biases, and particularly the “global bias”, had very little effect on array orientation with GDD, suggesting that the array becomes committed towards a particular orientation even before all information is fully integrated (Figure S3). Time traces of individual simulations on cylinders support this hypothesis. Figure 5, indeed, shows that, in most cases, *R*_2_ increased substantially after the a stable orientation was obtained (near horizontal part of the traces). At the times of rapid reorientation, on the other hand, *R*_2_ was relatively low. With both LDD microtubule-based and isotropic nucleation, the array orientation appeared more stable for transverse arrays than for longitudinal arrays, whereas for GDD, the stability looked more similar. Additionally, very few traces crossed the middle with GDD, whereas a full reorientation from longitudinal to transverse occurred in a substantial fraction of the runs with LDD and ISO.

One possible signature of less efficient whole cell information processing on larger geometries is the formation of local domains with competing orientations (see Figure 6 for some examples). To quantify how sensitive the different nucleation algorithms are to this, we compared simulations on square periodic domains of different sizes. With all three nucleation mechanisms, we found a reduction of the plateau *S*_2_ level of the aligned regime in (*G, S*_2_)-curves with increasing domain size (Figure S4). The plateau level was also lower for the elongated 20×80 *μm*^2^ than for the same surface area 40×40 *μm*^2^ domains, corroborating the idea that this is the result from local domain formation (Figure S4). With isotropic nucleation, the reduction was most pronounced, particularly on the largest (80×80 *μm*^2^) periodic squares. Plotting the same simulations as (*S*_2_, density) scatter plots (Figure 7) shows that local domain formation occurred most often and most extreme with isotropic nucleation, with some points with high density and high *G* (i.e., well inside the aligned regime) having (global) *S*_2_ values typical of disordered arrays. While these results offer a good explanation for the relative differences between LDD and ISO, they do not explain the insensitivity of GDD.

**Figure 6:**
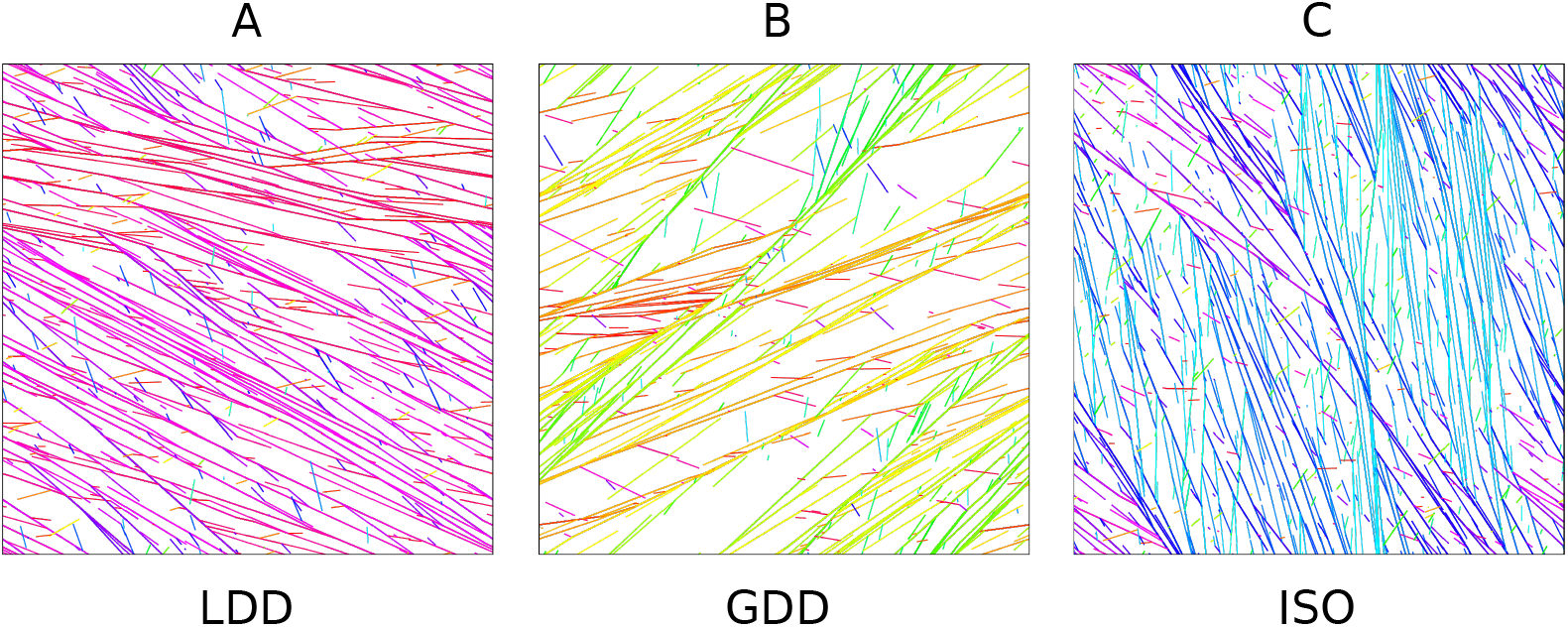
Local domain formation is much less pronounced in arrays obtained through LDD than through GDD and ISO nucleation modes. Snapshot at *T* = 3 × 10^4^ s (8 h, 20 min) of the microtubule array in a periodic square simulation domain of 80 × 80 *μ*m^2^ for (A) LDD: *S*_2_ = 0.90, *G* = −0.19, (B) GDD: *S*_2_ = 0.90, *G* = −0.19, and (C) ISO: *S*_2_ = 0.84, *G* = −0.14. Different colours corresponds to different orientations for a microtubule or a bundle: warm colours correspond to transverse and cool colours to longitudinal orientations.

**Figure 7:**
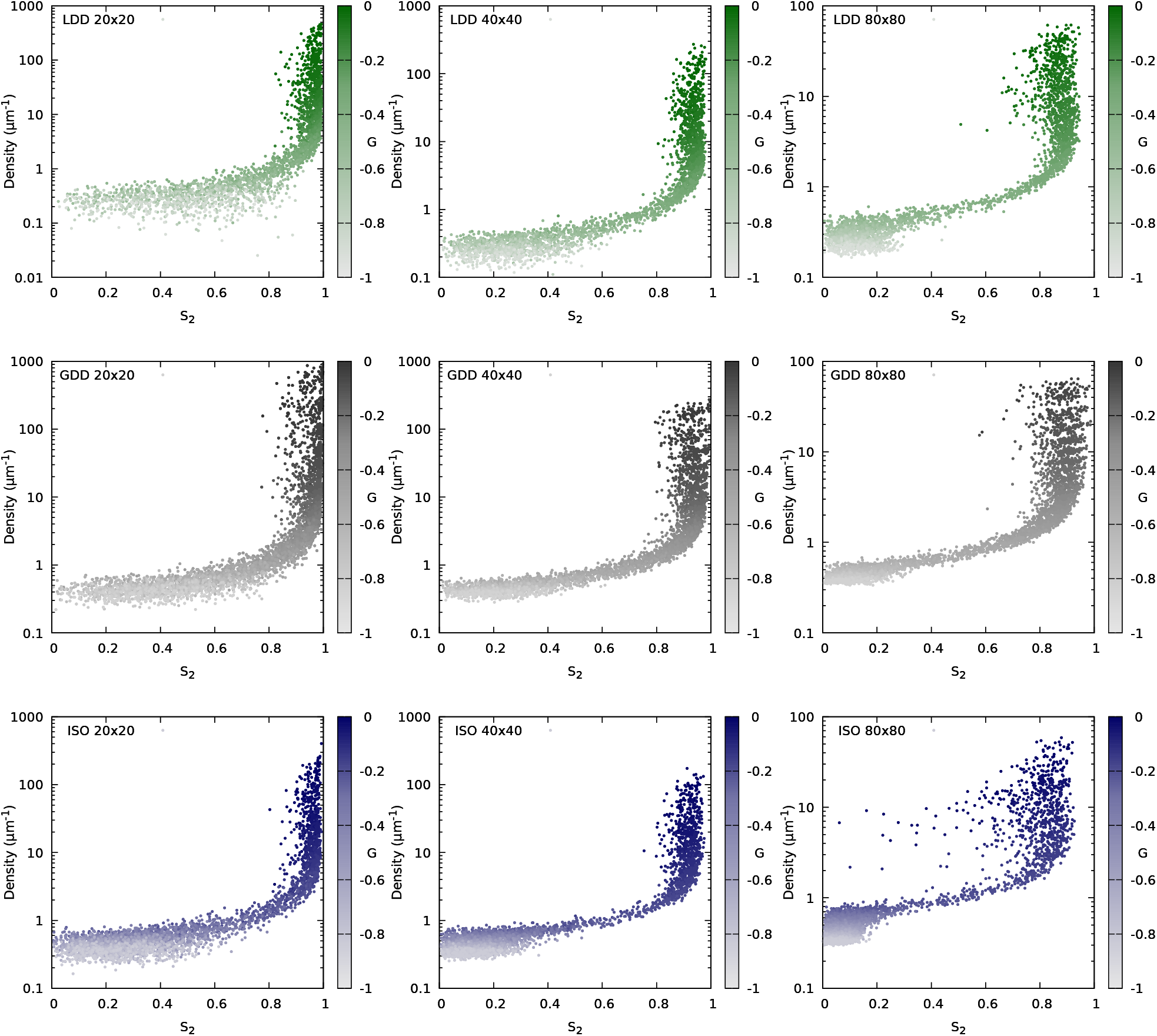
Clear signs of local domain formation with isotropic nucleation on large (80×80 *μ*m^2^ periodic) simulation domains. Scatter plots of density as a function of *S*_2_, coloured by *G* for the three nucleation modes (rows: LDD, GDD, ISO) and three domain sizes (columns: 20×20, 40×40 and 80×80 *μ*m^2^) as indicated in the graphs. Most points fall on a curve from low density and low *S*_2_ to high density and high *S*_2_ with increasing *G*. Points with relatively high *G* and density, but distinctly lower *S*_2_ than other points with similar *G* and density are in the regime of spontaneous alignment, but the occurrence of multiple aligned domains with different orientations reduces the global *S*_2_. The most cases, and most severe ones, occurred with isotropic nucleation on the largest domains. These plots are based on the same data as Figure S4, with 100 simulations per (target) *G* value. Simulation time: *T* = 3 × 10^4^ s (8 h, 20 min; lower for the highest *G* values for LDD and GDD, because not all simulations terminated in time).

#### 1.3.1 Cylinders and boxes have distinct impact on array orientation

To test whether the preference for transverse arose from the cylindrical shape or the aspect ratio of the geometry, we also investigated the preferred orientation on cylinders and boxes with different aspect ratios, while keeping the total surface area roughly constant (Figure 8, S5). On elongated boxes and cylinders, all three nucleation modes adopted either a transverse or a longitudinal orientation, albeit with different preferences for either. In particular, with LDD and ISO, the cylindrical geometry lead to a reduced preference for the longitudinal orientation compared to equally long boxes. With GDD, this did not happen. On square boxes, also a fully diagonal orientation appeared (all modes) and, with GDD, an orientation diagonal over two opposing faces and parallel along the two connecting edges. This orientation had a lower global degree of order (*R*_2_, see [11]) than other preferred orientations, probably the result of conflicts within the array^3^. We hypothesise that these internal conflicts have less impact when almost all array density is concentrated in a single closed band (expected for GDD) than with a broader, similarly oriented array and, therefore, these orientations are substantially less favourable with LDD and ISO (although they have been observed with ISO nucleation before, but with different parameters [20]). Further flattening the box lead to a strong preference for transverse orientations with all three nucleation modes, among which the orientations parallel to one face had the highest *R*_2_. The equivalent of the two types of diagonal orientations was only visible with GDD.

**Figure 8:**
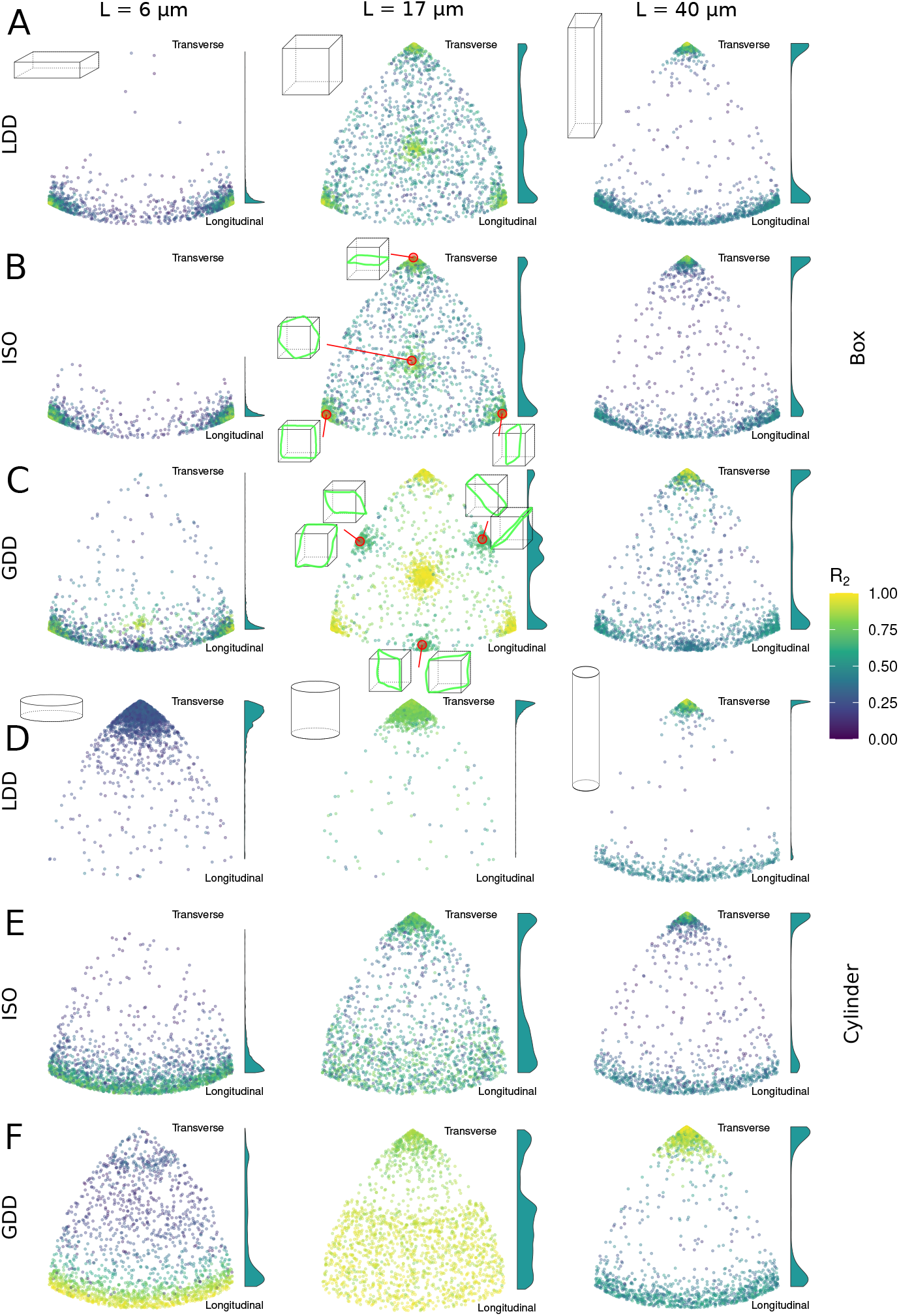
Nucleation mode has a strong impact on sensitivity to geometry. Orientation of *n* = 2000 arrays at *T* = 3 × 10^4^ s (8 h, 20 min) on boxes (A-C) and cylinders (D-F), all with the same total surface area. Individual points are coloured by *R*_2_ value. Note that this measure of alignment has an orientation and geometry dependent maximum value ≤ 1. The basis for the surface area is a cylinder of L=40 *μ*m and diameter of 12 *μ*m from figure 4A. For cylinders (D-F), L is varied as indicated, and diameter adjusted to maintain the same total surface area. This has approximately the same surface area as a LxWxW = 17x17x17 *μ*m^3^ box. For boxes (A-C), length L is varied as indicated and width W is adjusted accordingly to maintain the same total surface area. Cartoons are indicative of aspect ratios. Cartoons with green bands (ISO and GDD square box data) illustrate the array orientations belonging to the different positions in the plot. Histograms at the side of each plot show the relative distribution of transverse (top) to longitudinal (bottom) orientations. The highest peak in each histogram has a fixed height, i.e., the histograms are scaled differentially. In these simulations, *r*_*c*_ = 0.00175 s^−1^. For intermediate aspect ratios, see Figure S5.

Whereas the three nucleations modes showed relatively similar behaviour on boxes (Figure 8A-C), their behaviour on cylinders was very different (Figure 8D-F). With LDD, the transverse orientation was strongly favoured for most aspect ratios (Figure 8D, S5), albeit with increasing variation for flatter cylinders. With ISO (Figure 8E), transverse and roughly longitudinal were the most preferred orientations, and flattening the cylinder resulted in a gradual shift of preference from transverse to longitudinal. Just as increasing the nucleation rate of unbound complexes to the same level as microtubule-bound complexes made the alignment process of LDD more similar to ISO (Figure S2), increasing the unbound rate of LDD reduced the preference for transverse on the flattest cylinders and made the distribution of orientations more similar to ISO (Figure S6). With GDD (Figure 8F), there was a band of non-preferred oblique orientations, the width of which depended on the aspect ratio (Figure S5). Only on the flattest cylinder, GDD produced a strong preference for a longitudinal orientation. These results show that the impact of geometry itself is substantial even with isotropic nucleation. Microtubule-based nucleation further increases and sometimes alters this impact.

#### 1.3.2 Homogeneous and inhomogeneous arrays are sensitive to different aspects of cell geometry

GDD appeared almost insensitive to the cylindrical geometry, with very similar occurrences of longitudinal and transverse arrays. In contrast, on periodic (square) geometries, GDD was the only algorithm that showed a preference for certain orientations (Figure S7). These preferred orientations corresponded to the orientations of short closed paths that wrap around the geometry. This effect was most pronounced close to the transition to order (Figure S7D-F). Studying snapshots of individual simulations with such specific orientations showed that in these cases, there often was a single superbundle which wrapped back upon itself and, with time, absorbed most microtubule mass. With GDD, the parallel nucleations on the closed bundle increase its density, and, thereby, further increase the proportion of nucleations occuring from the closed bundle. With ISO, as well as with LDD nucleation, the additional density on the same bundle has no effect on the fraction of nucleations occurring from that bundle. So, only with GDD there is a positive feedback loop favouring closed superbundles and the corresponding array orientations.

On cylinders, the available closed paths are transverse and longitudinal. These coincide with the most adopted orientations with all nucleation modes (Figures 4, 5) Time traces of individual simulations on cylinders (Figure 5), nonetheless, show a difference in behaviour between GDD and the other two nucleation modes. With GDD microtubule-based nucleation, the array often adopted the closed path orientation closest to the initial array orientation, with almost no traces crossing the midline (± 45^°^ away from transverse). For snapshots of an example simulation, see Figure S8A. With the other two nucleation modes, however, multiple arrays (from n=50) were observed that reoriented from longitudinal to transverse (Figure 5A,B,E,F; for snapshots, see Figure S8B). Again, the deviating array behaviour with GDD could be explained from the strong positive feedback on closed bundles that occurs with GDD only: once a closed bundle is formed, the probability that a competing orientation established keeps decreasing with time, because the array outside the closed bundle is slowly drained.

## Discussion

In this work, we have developed a novel, computational time efficient nucleation algorithm for realistic microtubule-based nucleation that does not suffer from the inhomogeneity problem that haunted the field for over a decade [14, 15, 17, 32]. We showed that different nucleation mechanisms can have great impact on the overall behaviour of the cortical array. In particular, microtubule-based nucleation with local microtubule density-dependence increased the regime of spontaneous alignment and made the array more sensitive to subtle orientational cues from the cell’s geometry. Our simulations on different cell shapes and geometries demonstrate how the ability to realistically simulate microtubule-based nucleation opens up the possibility of quantitative comparison of different forces acting on array orientation.

While we believe that LDD is a much more realistic mechanism for nucleation than GDD, the formation of certain *inhomogeneous* array structures could benefit from GDD-like behaviour. For example, the preprophase band (PPB) somewhat resembles the single-banded arrays obtained with GDD (Figure 2F) [40]. To what extent would tuning between LDD-like and GDD-like nucleation be biochemically possible? From a theoretical perspective, the critical difference between the two nucleation-based algorithms that we discuss is whether the relative probability to attract a specific nucleation event to a particular small region linearly depends on the local microtubule density (GDD) or this probability saturates (LDD). The saturation is a natural effect of locally exploring nucleation complexes (e.g., diffusion after appearance at the membrane) and, therefore, to some extent unavoidable. The saturation may be delayed, however, by reducing the dwell time (→ smaller search area) of a nucleation complex at the membrane, or some co-factor that binds to the microtubule lattice and functions as a density sensor. Multiple factors, including Augmin, facilitate nucleation complex recruitment [31], which is observed to be biased towards microtubules [14], and local nucleation complex density increases proportionally as regular gaps start appearing in developing protoxylem [15]. Such effects make the nucleation a bit more GDD-like and indeed improve the translation of locally varying dynamic parameters to a pattern of distinct bands and gaps in simulated protoxylem arrays [14, 25]. In that context, array homogeneity at the whole-cell level is maintained, because appearance of nucleation complexes occurs throughout the cell [14, 25]. If the appearance of nucleation complexes would be biased towards a specific region via another patterning mechanism, this could, for example, support the formation of a “globally inhomogeneous” pattern like the PPB (see also [41]). Some experimental observations indeed support spatial variation in nucleation during PPB formation [40, 42]. To what extent the parameters that describe nucleation complex dynamics are varying over time and space and may be spatially regulated (e.g., via ROP proteins) remains a topic for future investigation.

Our simulations on different geometries show that cell shape can have a huge impact on the preferred set of orientations, stressing the importance of also considering realistic cell shapes [5,19,22]. In our simulations on cylinders, reliably establishing a longitudinal orientation *de novo* seemed hard, i.e., it required stronger differences in the orientation dependent rescue rate than the maximum we tested. A plausible explanation for the strong orienting effect is that the longitudinal orientation on a cylinder causes conflicts within the array, whereas the transverse orientation does not. Boxes have two specific longitudinal orientations without conflict, one or two of which disappear when the top and bottom face are distorted to fit into a cylindrical cell layer.

We found this strong bias even without a form of microtubule alignment with cell wall stresses (e.g., [22]) which, on a cylinder, are twice as large for transverse compared to longitudinal [43]. Taken together, these factors seem at odds with the natural requirement of producing longitudinal arrays when necessary. Extensive work on dark-grown hypocotyls, however, shows that reorientation from transverse to longitudinal can be a very active process in which, initially, the microtubules with the minority orientation are specifically amplified [23, 44–46]. A simpler model than the one used here, however, suggests that a relatively small bias, e.g., a 10% difference in growth speed, may be sufficient to subsequently tip the balance towards longitudinal, *when combined with a finite pool of total available tubulin* [23]. Note that this difference is smaller than our maximum 24% increase in rescue rate, whereas others have used a “modest” 80% difference in spontaneous catastrophe rate [47]. These differences among modellers strongly urge the experimental quantification of orientation dependent dynamics.

The orientation of the PPB also seems determined by an active process, that sometimes overrules the preceding interphase orientation, signified by the formation of a transient radial array structure [42]. This structure coincides with the observation of nuclear envelope-derived microtubules that join the cortex with independent orientation [42]. From a cortical perspective, this resembles a temporal increase of the “unbound” nucleation rate, with a spatially varying bias on orientation and intensity due to microtubule stiffness. This could aid in PPB positioning, by disrupting the established array structure in some places and strengthening it in others.

These active reorientation processes illustrate that a firm preference for one specific orientation can go hand in hand wider control over array orientation.

A major open question in the field is how plant cells weight different cues affecting the orientation of the cortical array. Such cues can be global, i.e., that always require information processing throughout the whole array, and local, i.e., that can be sensed (almost) everywhere in the cortex with similar intensity. Cell size, tuning the persistence length of microtubule growth [24, 25, 48], and the degree of parallel microtubule-based nucleation [17] can each shift the balance between local and global cues. The most prominent global cue derives directly from the cell geometry itself, as on many geometries, including cylinders, only a limited set of array orientations minimizes conflicts with itself [5, 20]. Cues like penalties for crossing (sharp) edges [5, 49] or cell faces with less favourable conditions can more selectively determine which orientations are realised [5, 20, 24]. In terms of information processing, these act as global cues.

In contrast, the co-alignment of microtubules with patterns of mechanical stress in the cell wall that seems to dominate in certain systems [21, 28, 38, 50–52], is typically assumed to happen via local cues, e.g., implemented as a slight bias in the microtubule growth direction towards the direction of (predicted) maximum wall stress [22], or increased growth propensity for co-aligned microtubules via the modulation of microtubule dynamics in simulation studies [24, 53]. The exact mechanism behind this observation remains elusive, but depends on katanin [38, 52]. This microtubule severing enzyme [54, 55] has a profound influence on alignment of the cortical array and is critically important for array reorientation in response to various internal or external cues [35, 38, 44, 52, 56]. Another potential local cue is the tendency of some microtubules to slowly reorient such that they minimize their curvature either globally or over a small region, e.g., favouring longitudinal orientation for non-interacting microtubules on an infinite cylinder [39, 57, 58]. Recent simulations show that this effect, indeed, competes with global geometry-based effects, including induced catastrophes at the edges of cylinders [47]. It would be interesting to continue this investigation using local density-dependent (LDD-like) nucleation, to test whether LDD favours the transverse orientation also if individual microtubule tips have some tendency to deflect towards longitudinal.

Our modelling advances enable the *in silico* investigation of the plant cortical array with a high degree of realism in a fast simulation environment. With our new algorithm, the simulation platform CorticalSim now is equipped with katanin severing [35], semiflexible microtubules [25], and more realistic, i.e., local density-dependent, microtubule-based nucleation that avoids the problem of artificial inhomogeneity. We have demonstrated that the mechanism of microtubule nucleation has profound impact on array orientation and other aspects of array behaviour. Moreover, there are many open questions that will benefit from fast and realistic simulation of microtubule-based nucleation.

## Methods

We simulate the cortical array with a stochastic platform that includes all microtubules as sets of connected line segments that interact. The platform, CorticalSim, is a fast, event-based microtubule simulation platform written in C++ [11]. In this platform, microtubules are confined to the surface of the cell membrane, in this manuscript modelled as closed cylinders or boxes, or a square/rectangular periodic domain. In the case of cylinders, microtubules have the same dynamic parameters on both the lateral surface and the two caps. In both the case of cylinders and boxes, we do not consider catastrophe associated with crossing sharp edges. Table 1 summarizes model parameters and their default values.

### Microtubule dynamics

The dynamics of microtubules comprises two main components: intrinsic dynamics and microtubule-microtubule interaction. We employed the Dogterom and Leibler model [60] to describe the intrinsic dynamics of individual microtubules augmented with hybrid treadmilling [59], see Figure 1C. Microtubules can be either in the growing state, where they extend at the plus-end with a growth speed of *v*^+^, or in the shrinking state, where they retract their plus-end with a shrinkage speed of *v*^−^. The minus-end of microtubules retracts at a treadmilling speed of *v*^tm^ independently of the state of the plus-end. Transitions between the growing and shrinking states, and *vice versa*, occur with catastrophe and rescue rates denoted as *r*_*c*_ and *r*_*r*_, respectively.

When a growing microtubule plus-end impinges on the lattice (the lateral, cylindrical surface of a microtubule, in this model treated as a segment) of another microtubule, the resulting type of interaction depends on the collision angle, comprised between 0^°^ and 90^°^. For collision angles smaller than *θ*_*c*_ = 40^°^, zippering occurs, causing the plus-end to reorient alongside the encountered microtubule. When the collision angle is wider than 40^°^, the impinging plus-end has a probability *P*_cat_ to undergo a collision-induced catastrophe, and a probability 1 − *P*_cat_ to create a crossover with the other microtubule, see Figure 1D. Microtubule bundles emerge as a result of the zippering process. We assume that the collision between a growing microtubule tip and a bundle is equivalent to a collision with a single microtubule. Likewise, when a microtubule within a bundle collides with another microtubule, it remains unaffected by the presence of other microtubules within the same bundle. CorticalSim can simulate microtubule severing events induced by the severing enzyme katanin [35]. However, for simplicity, we do not consider katanin severing in this manuscript.

### Microtubule nucleation

We explore three distinct modes of microtubule nucleation: isotropic nucleation (ISO), global density-dependent nucleation (GDD), and a novel local density-dependent (LDD) nucleation algorithm. The latter consists of a new computational approach, and we compare its performance against the former two methods, already well-studied in the literature [11, 17, 32, 33]. Regardless of the chosen nucleation mode, we assume that nucleation complex availability and behaviour are constant over time. The LDD algorithm incorporates the following experimental observations: 1) nucleation primarily occurs from the lattice of existing microtubules [12,13,30], 2) with a distribution of relative nucleation angles based on [12], and 3) observations of sparse, oryzalin treated, arrays show that nucleation complexes can appear in empty parts of the array, but have a higher nucleation efficiency after association with a microtubule [13, 14].

#### Isotropic (ISO)

In the case of isotropic nucleation, new microtubules are nucleated at a rate *r*_*n*_. The location and orientation are randomly selected with uniform probability.

#### Global density-dependence (GDD)

In the global density-dependent nucleation (GDD) case, which was developed in [17], nucleation can be either microtubule-based or background unbound nucleation. Given the total density of tubulin *ρ* used by microtubules, the rate at which microtubule-based nucleation occurs is

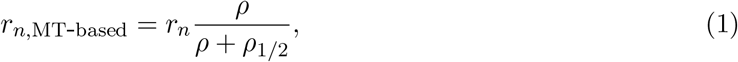

where *ρ*_1*/*2_ is an equilibrium constant determining the density at which half of the nucleations are microtubule-based [18]. Consequently, isotropic background nucleation occurs at a rate *r*_*n*,unbound_ = (*r*_*n*_ − *r*_*n*,MT-based_). This approach ensures that initial nucleations at the microtubule-free membrane are unbound, while a small fraction of unbound nucleations occurs even at higher densities. In our simulations, the locations of microtubule-based nucleation events are uniformly distributed along the length of all available microtubules.

We modelled the angular distribution of microtubule-based nucleations as proposed by Deinum et al. [17]. Their nucleation mode includes three components: forward (towards the plus-end of the parent microtubule) with probability *f*_forward_, backward (towards the minus-end of the parent microtubule) with probability *f*_backward_, and the remaining probability sideways, split between left (*f*_left_) and right (*f*_right_). The sideways angles are modelled proportional to the relative partial surface area from a focal point of an ellipse with eccentricity *ϵ*, with its main axis oriented with branching angle *θ*_*b*_ relative to the parent model. Parameters *ϵ* and *θ*_*b*_ are based on the data by Chan and Lloyd [12]. The full nucleation distribution function, denoted as *ν*(*θ*), where *θ* is the angle between the direction of the parent and the daughter microtubules, combines these components:

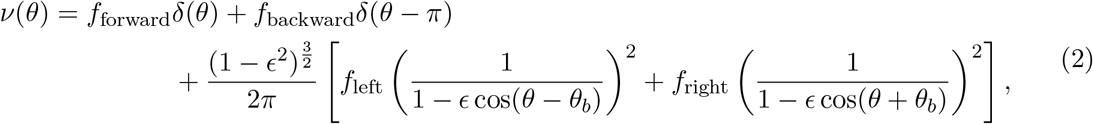

where *δ* is the Dirac *δ*-distribution.

#### Local density-dependence (LDD)

New nucleation complexes appear at the membrane at rate *r*_ins_ following the observation of nucleation complexes freely diffusing at the membrane [14]. These complexes are virtually placed at a random position **x**_0_ within the simulation domain. For convenience, and without loss of generality, we set **x**_0_ = (0, 0) in the remainder of this section. To define a region for nucleation, we consider a circle with a radius *R* centered at **x**_0_, which we call the nucleation area. Within this area, we draw *N* radii, each referred to as a meta-trajectory. The first meta-trajectory is drawn at a random angle *θ*_1_ ∈ [0, 2*π*) in the simulation domain, while subsequent meta-trajectories have angles *θ*_*i*_ = *θ*_*i*−1_ + 2*π/N*, see Figure 9A and B. A meta-trajectory labelled by *i* may either intersect the lattice of a microtubule at a distance *d*_*i*_ from the centre or reach the boundary of the nucleation area, with *d*_*i*_ being equal to the radius *R* in the latter case. When a meta-trajectory intersects a microtubule lattice, it stops there (Figure 9B). We approximate the diffusion process by assuming that the nucleation complex can either reach these intersections by diffusion or diffuse away from the nucleation area. Here, we use *N* = 6 to reasonably represent the possible directions of diffusion of nucleation complexes.

**Figure 9:**
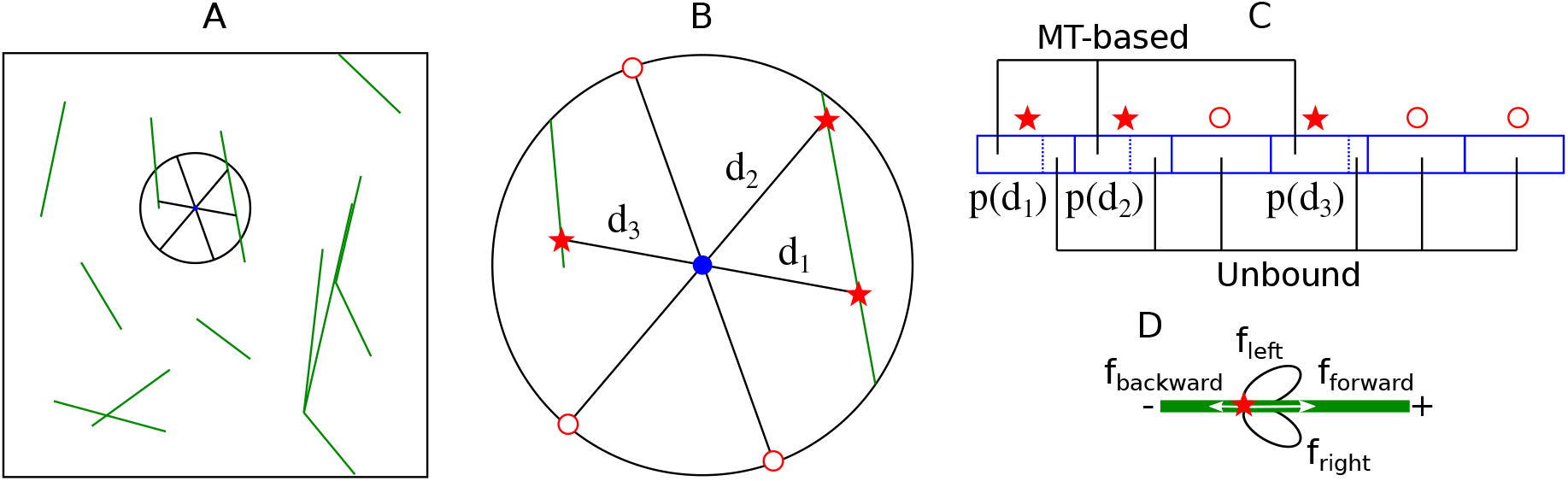
Schematic overview of the LDD nucleation algorithm. (A) The position in the simulation domain where a nucleation complex (tentatively) appears is selected randomly, with uniform distribution. (B) Such a position, represented by the blue dot, is the centre of the nucleation area (the area within in the black circle with exploration radius *R*). *N* meta-trajectories are drawn equidistant from one another and with random orientation. Meta-trajectories are drawn up until the boundary of the nucleation area or the lattice of the first intersecting microtubule, whichever is closer. Whether the resulting nucleation is microtubule-based or unbound and, in the former case, the parent microtubule and the nucleation location, is decided stochastically with probability *p*(*d*_*i*_) weighted over the length of meta-trajectories, according to Equation (4). (D) In a microtubule-based nucleation, the angles follow Equation (2) [17]: a newly-nucleated microtubule initially grows along the parent microtubule with probability *f*_forward_, in the opposite direction with probability *f*_backward_, or branches to either side with probability *f*_left*/*right_. The branching angle is determined according to a distribution that closely matches the data in [12] (Equation (2)).

Following the appearance of a nucleation complex at the membrane, three outcomes are possible: (i) the nucleation complex dissociates, (ii) an unbound nucleation occurs, (iii) or the complex reaches the lattice of a microtubule. After reaching a microtubule lattice (case iii), the complex may either produce a microtubule-based nucleation or dissociate before that. In experiments, the overall probability that a nucleation complex dissociated without nucleation depended on whether such a complex was freely diffusing or attached to a microtubule lattice. In the former case, the probability of dissociation is 98%, in the latter 76% [13, 14]. Ideally, the nucleation process should be modelled as a two-stage mechanism, with dissociation probabilities depending on time until reaching a microtubule lattice. Unfortunately, however, this cannot be parametrized with existing experimental data and performing the relevant measurements would be very challenging, because of the high density of most cortical arrays [13, 14]. We, therefore, adopt a pragmatic approach. We first determine the likelihood of a complex reaching a microtubule lattice, leading to a tentative microtubule-based or unbound nucleation event. Subsequently, part of the tentative nucleation events are rejected based on the above dissociation probabilities.

We assume a nucleation complex tentatively reaches a microtubule lattice if it does not produce an unbound nucleation before reaching the site via a 2D diffusion process. For this, we assume a constant unbound nucleation rate *r*_*u*_, and take the time required to reach the site as the expected time for a mean squared displacement of distance *d*_*i*_:

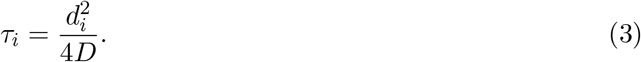

Consequently, the probability *p* that a microtubule tentatively reaches the lattice intersected by the meta-trajectory *i* before unbound nucleation is given by:

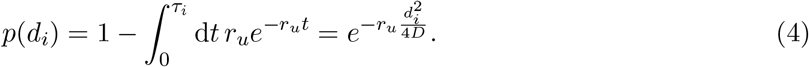

Hence, 0.24*p*(*d*_*i*_) is the probability of a microtubule-based nucleation on meta-trajectory *i*. Similarly, 0.02 (1 − *p*(*d*_*i*_)) is the probability of unbound nucleation on meta-trajectory *i*. In our simulations, we used a maximum meta-trajectory length of *R* = 1.5*μ*m because, given our choice of parameters values (Table 1), the probability *p*(*R*) of a nucleation complex intersecting with a microtubule lattice prior to nucleation becomes negligible beyond this length. Based on this, the probability of microtubule-based nucleation after a nucleation complex appears at the membrane is

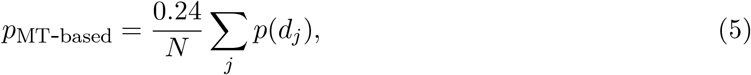

where the sum over *j* only runs over meta-trajectories that intersect a microtubule, while the probability of unbound nucleation is 0.02(1 − ∑ _*j*_ *p*(*d*_*j*_)*/N*), see Figure 9C. Once the method was well established, we further improved computational efficiency using an equivalent, more efficient parametrization. This means a reduced appearance rate 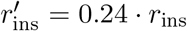, accepting all tentative bound nucleations, and accepting tentative unbound nucleations with probability 0.08333 = 0.02/0.24. Both parametrizations are used in the current figures.

Verification simulations with LDD on a planar simulation domain show how the fraction of microtubule-based nucleation increases with increasing microtubule density and the nucleation rate increases to equilibrium levels (Figure S10). At a high density around 5 *μ*m^−1^, the fraction microtubule-based nucleation was somewhat lower than reported before (95% when using local density, Figure S10, compared to 98% in [13]), which can likely be explained by the fact that, here, we do not take into account that the appearance of nucleation complexes at the membrane is biased towards existing microtubules [13, 14]. As expected, the observed relation between microtubule density and fraction of microtubule-based nucleation was tighter when using the local microtubule concentration than the global one (Figure S10).

### Degree of alignment and orientation of the array

To assess the alignment of microtubules in a planar system, we use a nematic order parameter *S*_2_ that accounts for the contribution of individual microtubule segments proportionally weighted on their length [17]. For planar geometries, *S*_2_ is defined by

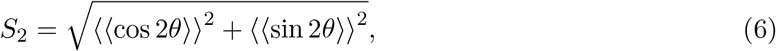

where the double brackets denote a length-weighted average over all microtubule segments. *S*_2_ has a value of 0 for an isotropic system and a value of 1 for a system where microtubules are perfectly aligned. For non-planar geometries, we use the order parameter *R*_2_ ∈ [0, 1] which extends the definition of nematic order parameter to surfaces in the 3D space as defined in [11]. As the standard extension of the *S*_2_ order parameter to 3D would produce non-zero values for isotropic arrays on cylinders and boxes, because of the confinement of the microtubules to the surface, the definition of *R*_2_ includes a correction for the expected value for a fully isotropic array. Consequently, the maximum *R*_2_ value depends on array orientation: for example, it is 1 for transverse arrays and (1+D/L)/2 for longitudinal arrays on cylindrical arrays with length L *>* diameter D. The minimal value is always 0. Only in Figure 5, *R*_2_ values are renormalized by this maximum.

We couple the alignment order parameter *S*_2_ with the overall array orientation Θ_2_:

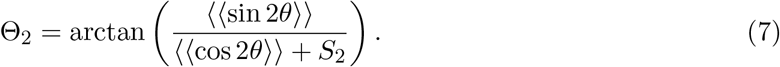

On non-planar geometries (capped cylinders and boxes in this manuscript), we use the definition of orientation vector Θ_2_ coupled to *R*_2_ [11]. This is the angle between the cylinder axis and the direction most *avoided* by the array, i.e., the predicted preferred direction of cell expansion. Note that this is the opposite of *S*_2_-based Θ_2_. We, therefore, indicate the longitudinal and transverse orientation alongside the relevant Figures to avoid confusion.

#### Navigating parameter space with control parameter *G*

The driver of spontaneous alignment in the cortical array is the occurrence of sufficient interactions within the average microtubule length. This is quantified by a single control parameter *G* [8, 9], which is the ratio of two length scales: an interaction length scale (left factor) and the average length of non-interacting microtubules (right factor). We use a version of *G* adjusted for minus end treadmilling [17]:

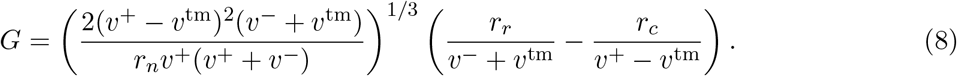

Note that we restrict ourselves to *G <* 0, i.e., when microtubules are in the so-called bounded-growth regime, where they have a finite life span and finite intrinsic length. In this case, spontaneous alignment occurs if *G > G*^∗^, some threshold that depends on the details of microtubule-microtubule interactions.

For ISO and GDD nucleation modes, the nucleation rate *r*_*n*_ is an explicit simulation parameter. For LDD nucleation, however, *r*_*n*_ results from the interplay between the rate at which nucleation complexes appear at the membrane *r*_ins_ and rejection of nucleations, which in turn depends on the density and distribution of the microtubules. Therefore, we calculate the effective nucleation rate to be used for *G* in Equation (8) as

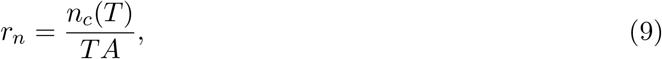

where *n*_*c*_(*T*) is the number of nucleation events occurred until time *T*, and *A* is the area of the simulation domain. In principle, *n*_*c*_ might be non-linear in *T*, making our proposed computation of *r*_*n*_ overly-simplistic. However, our test showed that *n*_*c*_(*T*) is approximately linear in *T*, for *T <* 3 × 10^4^ s (8 h, 20 min), see Figure S9.

#### Global and local bias on cylindrical geometry

We incorporated two distinct directional biases independently: a global bias, represented by an 8% increase in the catastrophe rate *r*_*c*_ at the cylinder caps, and a local bias, with a maximum *b*_max_ (default: 8%) orientation dependent increase of the rescue rate *r*_*r*_ for individual microtubules. Similar to others [23, 47, 61], we used a cosine function to model the increase:

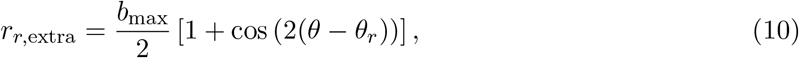

where *θ* denotes the orientation of the microtubule plus-end, *θ*_*r*_ represents the angle at which the rescue rate is maximum, and *b*_max_ is the maximal increase factor when *θ* = *θ*_*r*_.

## Supplementary Figures

**Figure S1:**
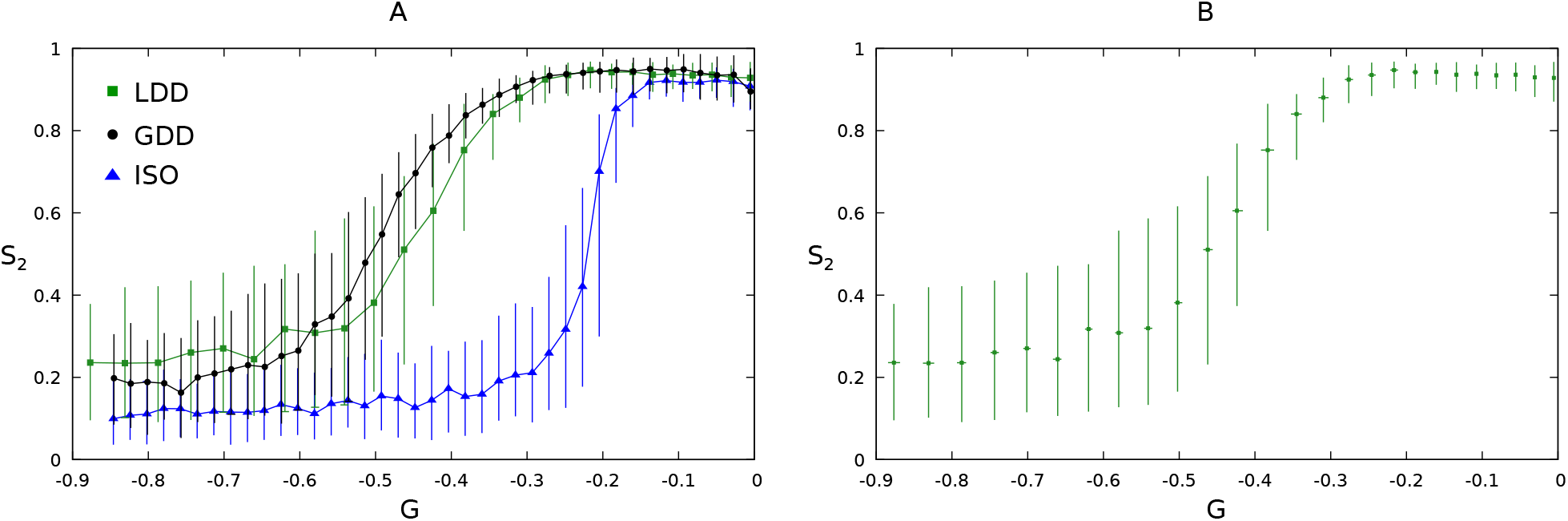
(A) Median over 100 independent simulation runs of the order parameter *S*_2_ as function of the control parameter *G* at *T* = 3 × 10^4^ s (8 h, 20 min), for LDD, GDD, and ISO nucleation modes in a squared geometry with sizes *L* = 40*μ*m and *H* = 40*μ*m, as depicted in Figure 3. Error bars represent the 10th and 90th percentile. (B) Median of the parameter *S*_2_ as a function of *G* for LDD as in panel (A), with the addition of error bars on the *G*-axis.

**Figure S2:**
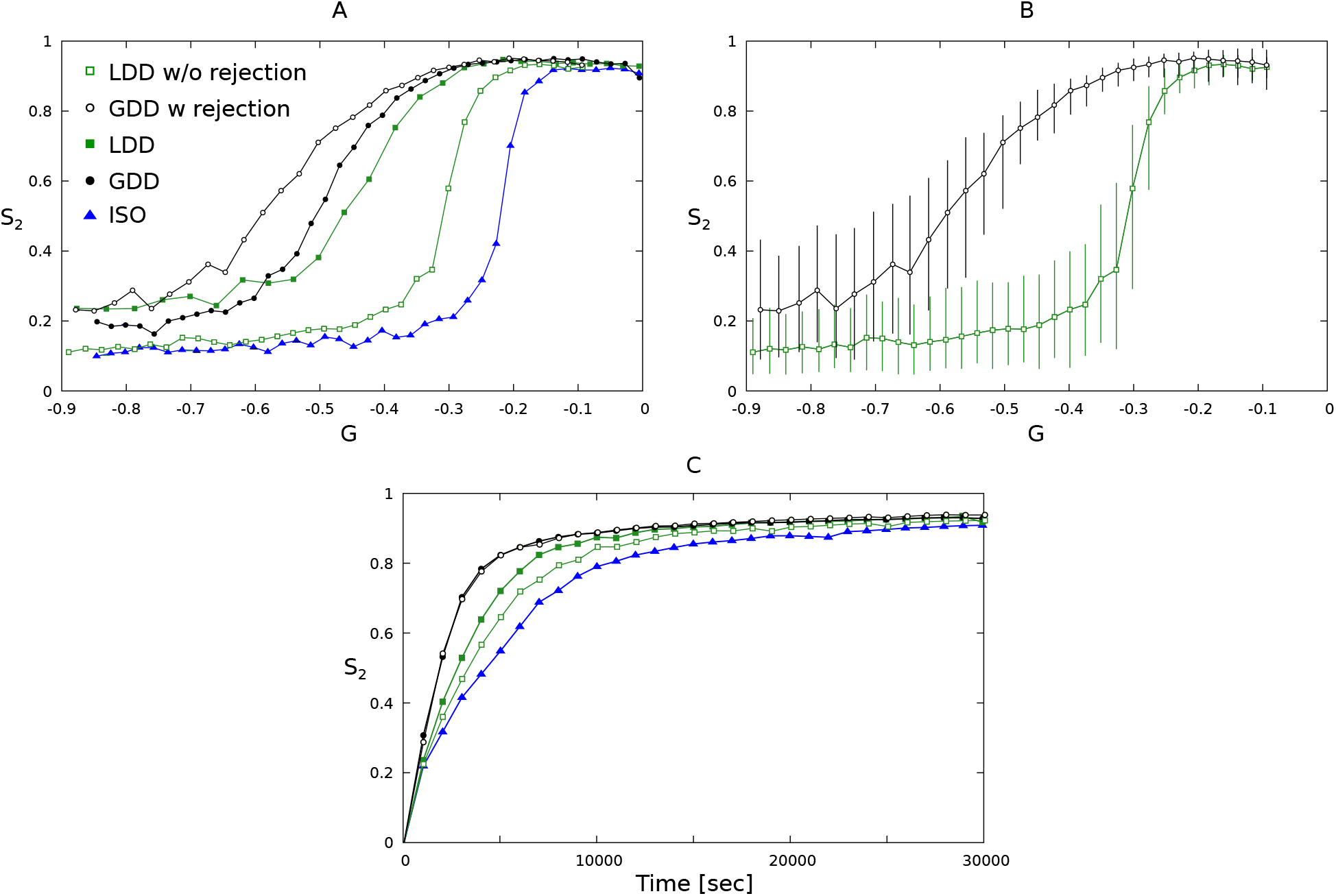
Median over 100 independent simulation runs of the order parameter *S*_2_ (A,B) as function of the control parameter *G* at *T* = 3 × 10^4^ s (8 h, 20 min), and (C) as a function of time for LDD, GDD, and ISO nucleation modes as in Figures 3 and S1, supplemented by LDD without rejection of unbound nucleations (i.e., same rate on and off microtubules), and GDD with rejection of 91.7% of unbound nucleation (i.e., off-microtubule rate reduced), on a square periodic geometry with sizes *L* = 40*μ*m and *H* = 40*μ*m. (C) In these simulations, in the case of LDD without rejection *r*_*c*_ = 0.0025 s^-1^, in the case of GDD with rejection *r*_*c*_ = 0.00325 s^-1^. Error bars represent the 10th and 90th percentile.

**Figure S3:**
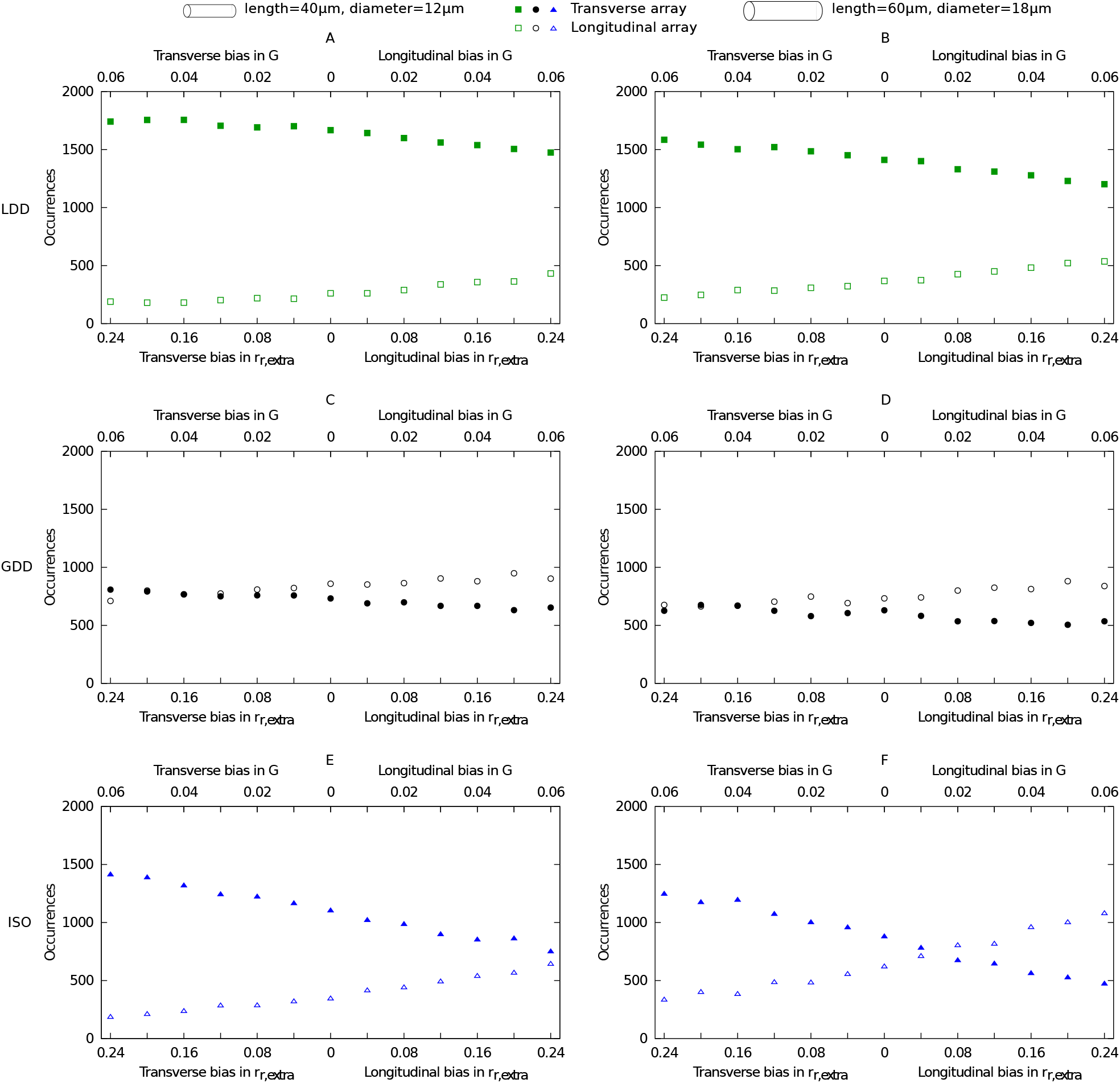
LDD selects for more transverse arrays than GDD and ISO. Number of aligned arrays at *T* = 3 × 10^4^ s (8 h, 20 min) with a transverse (filled symbols) or longitudinal (empty symbols) for the LDD (green squares), GDD (black circles), and ISO (blue triangles) cases, for different values of the longitudinal bias in *r*_*r*,extra_ according to Equation (10). The simulation domain is a cylinder of (A) 40*μ*m length and 12*μ*m diameter, and (B) 60*μ*m length and 18*μ*m diameter. The upper horizontal axis translates the increased rescue rate into the maximum increase in *G* in either the transverse or longitudinal directions. Error bars have been omitted because they are approximately the same size as the symbols. In these simulations, *r*_*c*_ = 0.00225 s^−1^.

**Figure S4:**
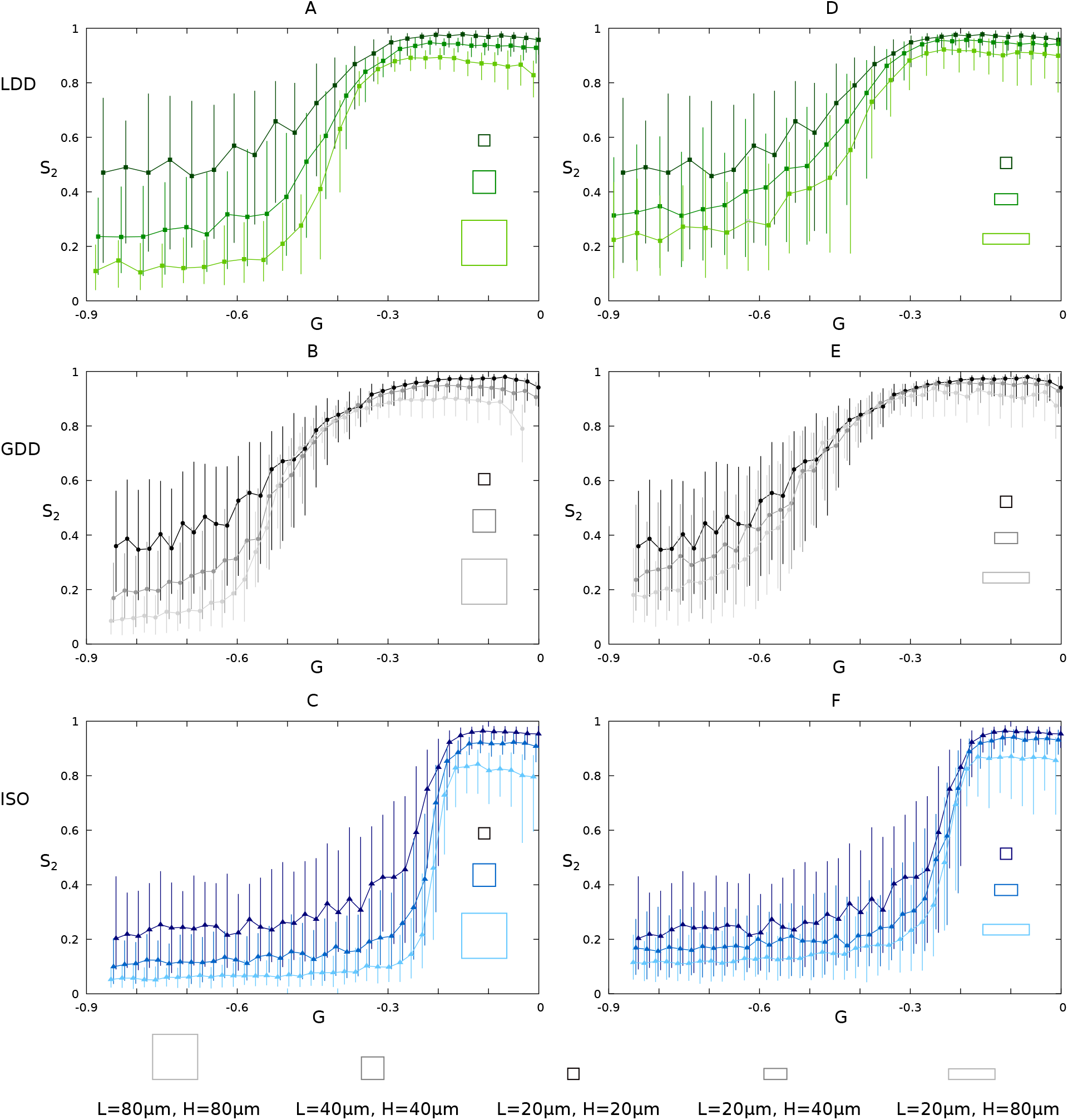
Dependence of global alignment on geometry size and aspect ratio for all nucleation modes. Median over 100 simulation runs of the degree of alignment *S*_2_ as a function of the control parameter *G* for LDD, GDD, and ISO nucleation modes for (A-C) increasing size and (D-F) increasing height-to-length aspect ratio of the planar periodic simulation domain, at *T* = 3 × 10^4^ s (8 h, 20 min). (B,C,E,F) In the ISO and GDD cases, results for the *L* = 20*μ*m, *H* = 20*μ*m grid are shifted to the right on the *G* axis by 0.005, whereas results for the *L* = 80*μ*m, *H* = 80*μ*m and the *L* = 20*μ*m, *H* = 80*μ*m grids are shifted to the left by 0.005 for visibility of the error bars. Error bars represent the 10^th^ to 90^th^ percentile.

**Figure S5:**
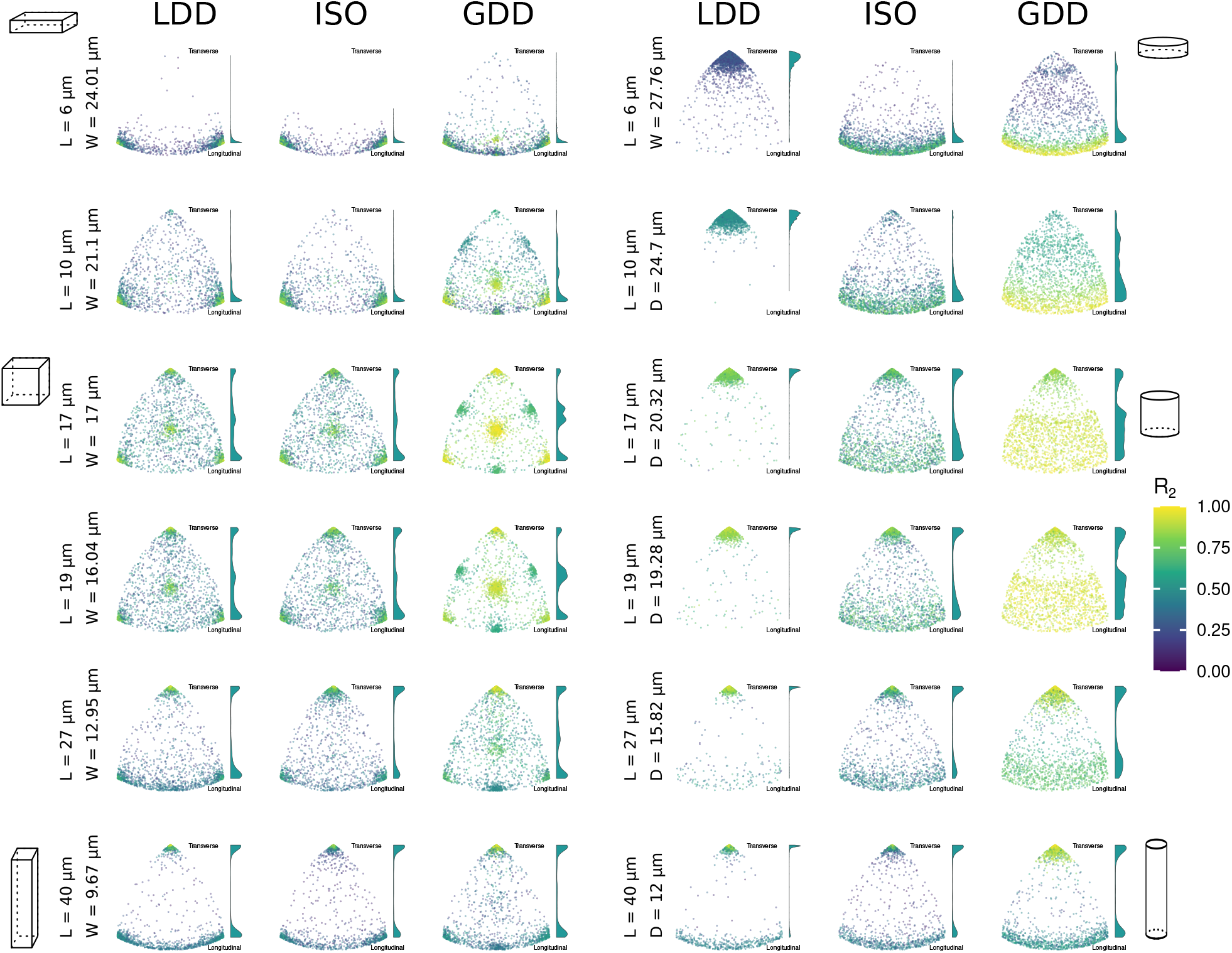
Extension of figure 8: Nucleation mode has a strong impact on sensitivity to geometry. Orientation of *n* = 2000 arrays at *T* = 3 × 10^4^ s (8 h, 20 min) on different geometries with the same total surface area. Individual points are coloured by *R*_2_ value. Note that this measure of alignment has an orientation and geometry dependent maximum value ≤ 1. The basis for the surface area is a cylinder of L = 40 *μ*m and diameter D = 2 *μ*m from figure 4A. For cylinders, L is varied as indicated, and diameter adjusted to maintain the same total surface area. This has approximately the same surface area as a LxWxW = 17x17x17 *μ*m^3^ box. For boxes, length L is varied as indicated and width W is adjusted accordingly to maintain the same total surface area. Cartoons are indicative of aspect ratios. Cartoons with green bands (ISO and GDD square box data) illustrate the array orientations belonging to the different positions in the plot. Histograms at the side of each plot show the relative distribution of transverse (top) to longitudinal (bottom) orientations. The highest peak in each histogram has a fixed height, i.e., the histograms are scale differentially. In these simulations, *r*_*c*_ = 0.00175 s^−1^.

**Figure S6:**
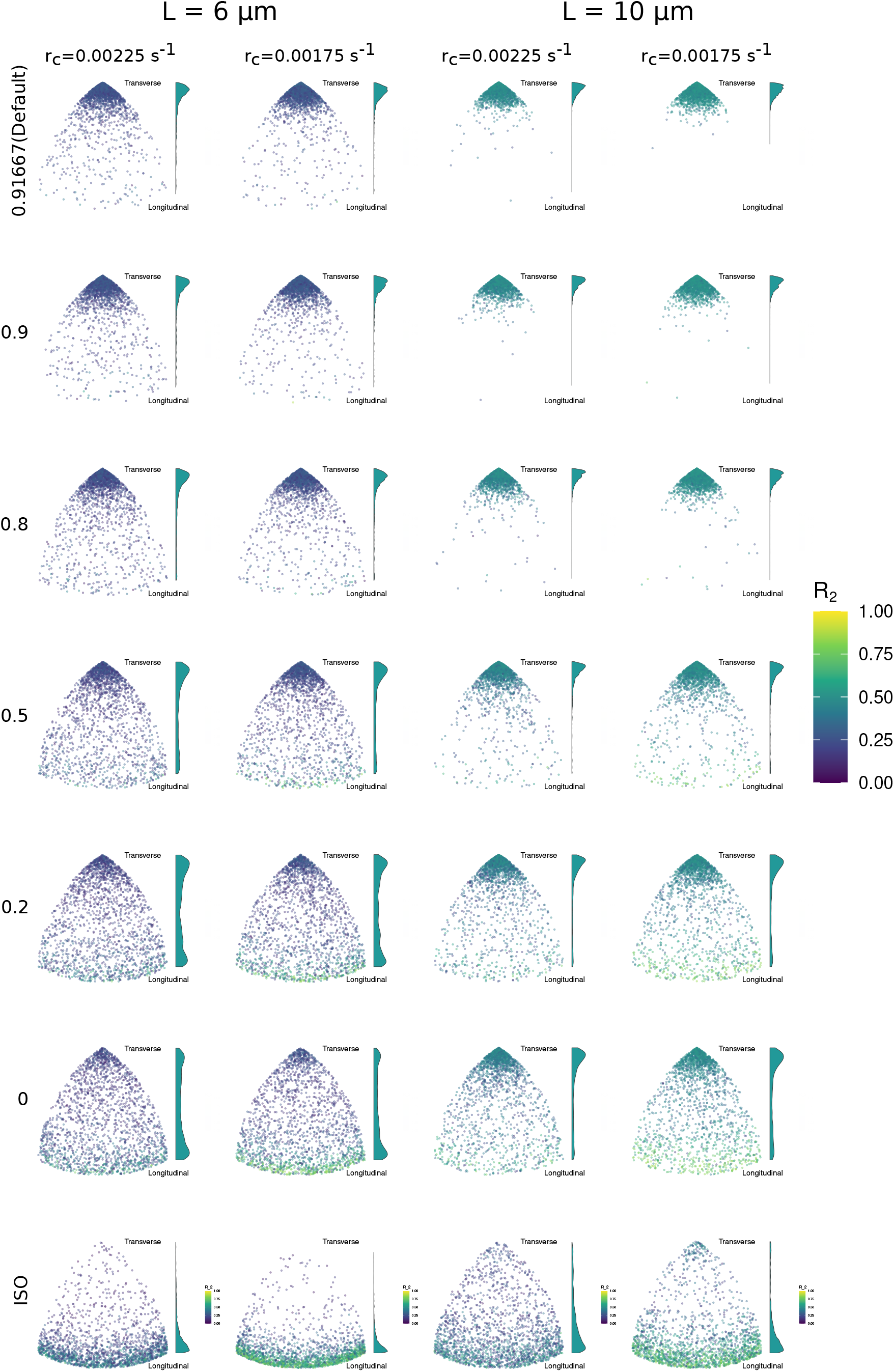
(Relative) rejection probability of unbound nucleations has a strong impact on array orientation with LDD nucleation. Orientation of *n* = 2000 arrays at *T* = 3 × 10^4^ s (8 h, 20 min) on different geometries with the same total surface area as in figure 8. From top to bottom, the rejection probability of unbound nucleations decreases as indicated. No bound nucleations are rejected, so the acceptance ratio is 1 − *P*_*reject*,*unbound*_. The bottom row shows isotropic nucleation as a reference. As the nucleation rates for bound and unbound complexes become more equal (=towards the bottom), LDD starts behaving more similar to ISO, but always retains a stronger preference for transverse than ISO. Individual points are coloured by *R*_2_ value. Note that this measure of alignment has an orientation and geometry dependent maximum value ≤ 1. The basis for the surface area is a cylinder of L=40 *μ*m and diameter of 12 *μ*m from figure 4A. For cylinders, L is varied as indicated, and diameter adjusted to maintain the same total surface area. Histograms at the side of each plot show the relative distribution of transverse (top) to longitudinal (bottom) orientations. The highest peak in each histogram has a fixed height, i.e., the histograms are scale differentially. In these simulations, *r*_*c*_ = 0.00225 s^−1^ or *r*_*c*_ = 0.00175 s^−1^, as indicated.

**Figure S7:**
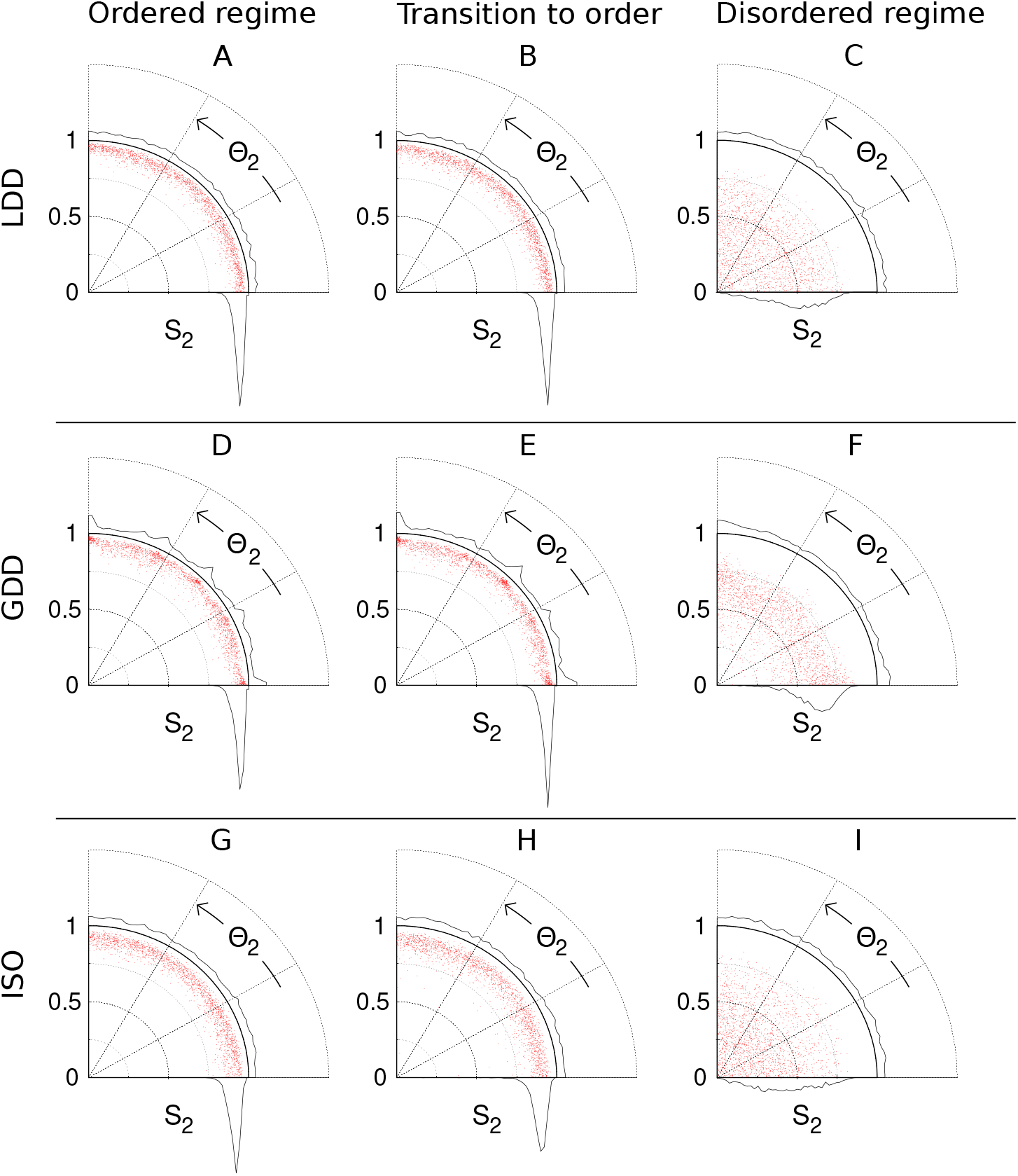
LDD nucleation is bias-free for orientation on a square periodic geometry, suggesting that the preference for the shortest few closed paths around this geometry with GDD is a consequence of the inhomogeneity problem. Distribution of *S*_2_ and Θ_2_ for three different regimes . Each panel contains the result in terms of alignment *S*_2_ and orientation Θ_2_ of 2000 independent runs at *T* = 3 × 10^4^ s (8 h, 20 min). A point located along the horizontal axis represents a perfectly transverse array orientation, while a point on the vertical axis represents a perfectly longitudinal orientation. The distance of the point from the origin indicates the strength of the alignment, with greater distances indicating a higher degree of alignment. The data points are collected to generate histograms for the distribution of *S*_2_ (horizontal axis) and Θ_2_ (vertical axis). *G* values: (A) *G* = −0.22, (B) *G* = −0.25, (C) *G* = −0.42, (D) *G* = −0.23, (E) *G* = −0.25, (F) *G* = −0.54, (G) *G* = −0.16, (H) *G* = −0.18, (I) *G* = −0.25. Panels (A), (D), and (G) correspond to the empty markers of Figure 3A.

**Figure S8:**
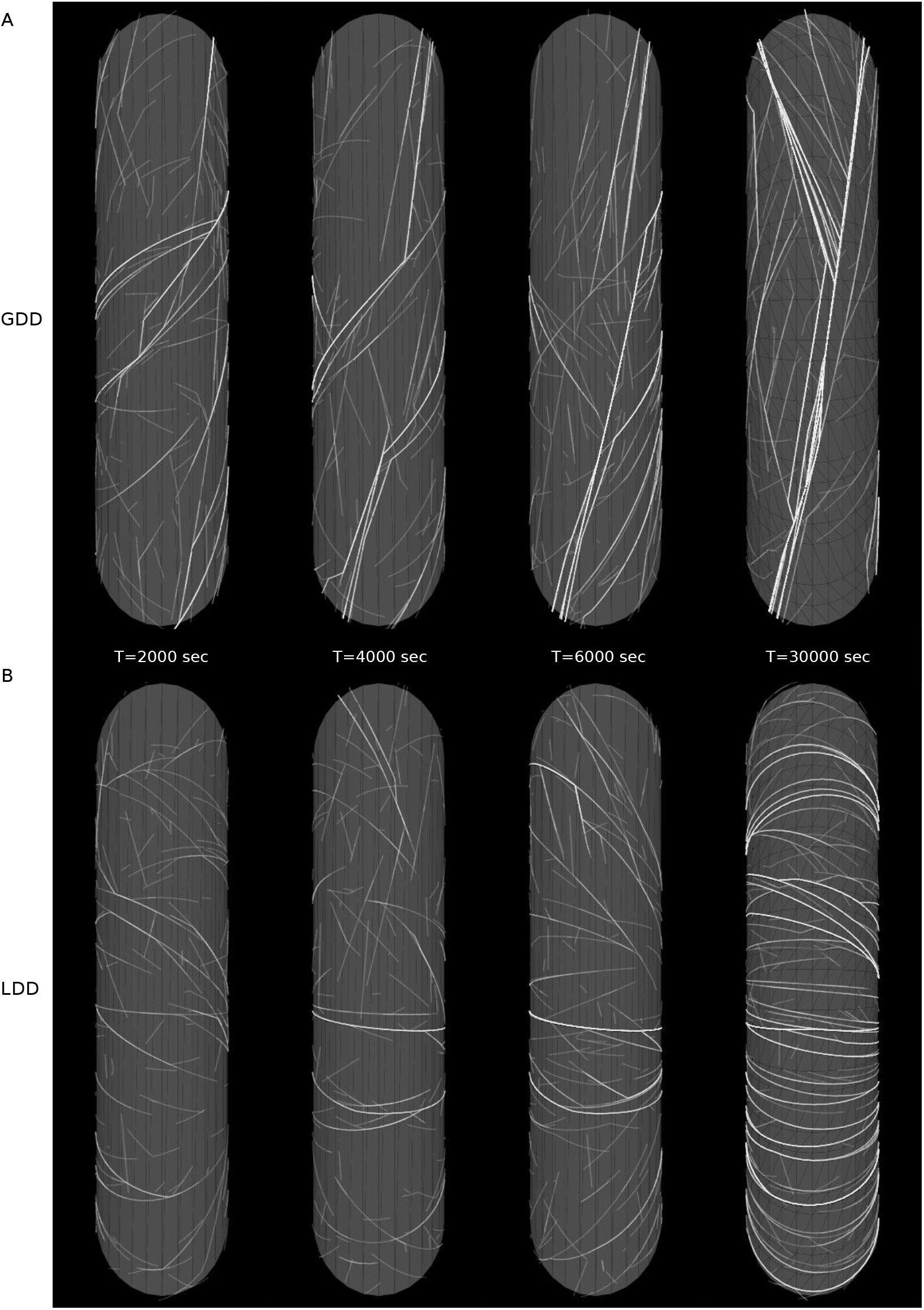
Snapshot of the simulated cortical array for (A) LDD and (B) GDD nucleation modes at four different time stamps. Simulation domain: cylinder with length= 40*μ*m, diameter= 12*μ*m. In these simulations: *r*_*c*_ = 0.0025 s^-1^.

**Figure S9:**
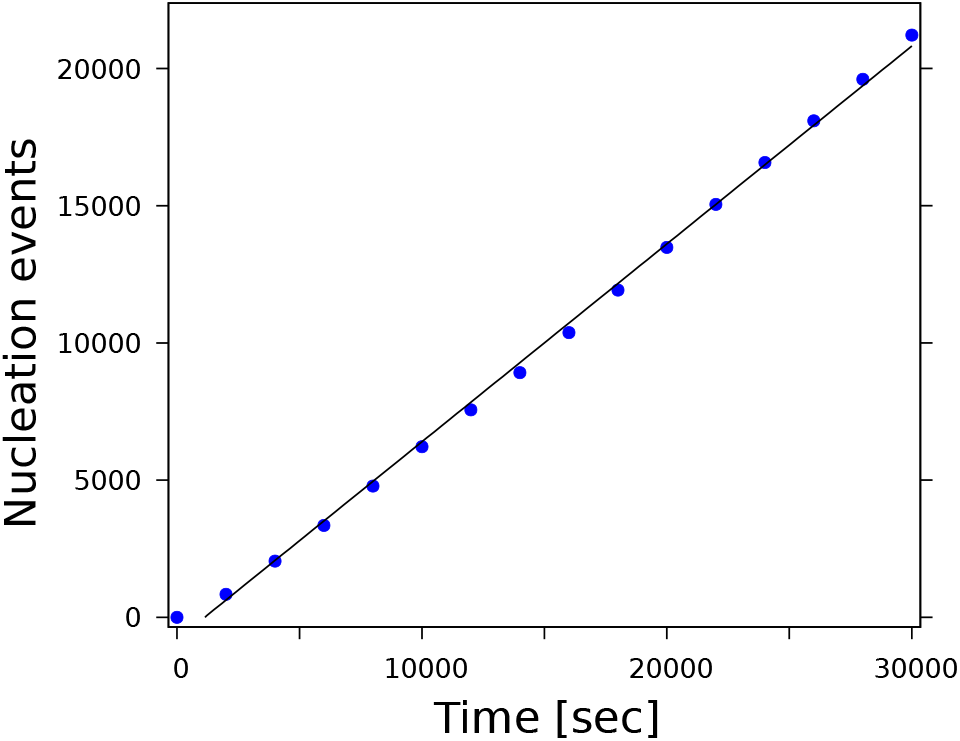
The nucleation rate remains roughly constant throughout the simulation. Cumulative number of nucleation events as a function of time for one simulation run with LDD nucleation mode and in a square simulation domain with periodic boundary conditions and size *L* = 40*μ*m. The black line corresponds to the fitting function *f* (*t*) = *a* + *bt*, with *a* = −809 ± 153 and *b* = 0.72 ± 0.01. In this simulation, *r*_*c*_ = 0.0025 s^-1^.

**Figure S10:**
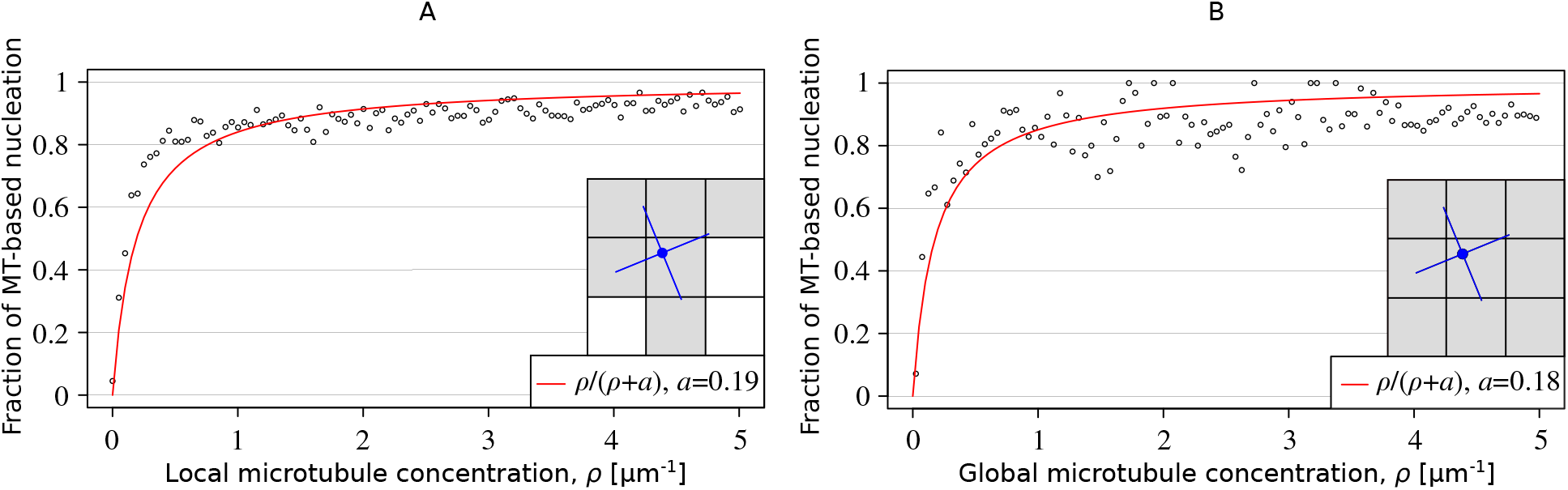
LDD microtubule-based nucleation naturally results in a local microtubule density-dependent fraction of microtubule-based nucleation. Fraction of microtubule-based nucleation as a function of (A) local and (B) global microtubule density (*ρ*) during a single simulation run. In this simulation, *r*_*c*_ = 0.0025 s^-1^. The global microtubule density (B) was computed as expected as *ρ* = ∑ _*i*_ *l*_*i*_*/A*, where *l*_*i*_ represents the length of the *i*-th microtubule, the summation runs over all microtubules, and *A* denotes the area of the simulation domain. The local microtubule density (A) was determined via 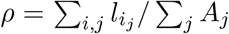. In this expression, 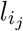 represents the length of the segment of the *i*-th microtubule within region *j* with area *A*_*j*_, and *j* runs over all regions touched by the metatrajectories drawn from the appearance point of a nucleation complex, as shown by the schematic in panel A. The curves fit through the data are of the same shape as the formula used for determining the fraction of microtubule-based nucleation with the GDD microtubule-based nucleation algorithm.

## Footnotes

1 ”CorticalSimple”, the software employed by Jacobs et al. [14], gives all microtubules an exactly transverse orientation, so there are no collisional interactions.

2 Due to the vagaries of the publication process, we have already published one research paper that uses the new nucleation algorithm in CorticalSim, in the specific context xylem patterning [25], which cites a preprint of this work for nucleation. Here, we describe the algorithm itself and investigate fundamental aspects of its behaviour and impact.

3 Note that *R*_2_ contains a correction for the isotropic expectation, so for the flat and elongated shapes, the maximum attainable *R*_2_ value for longitudinal *versus* transverse orientation can be very different. A difference in *R*_2_ for similar orientations, however, remains meaningful.

